# Myosin II-FAK-paxillin Signaling Governs Mechanosensitive Responses of Neuronal Cells to Magnetic Stimulation

**DOI:** 10.64898/2026.11.19.766553

**Authors:** Md Iqbal Kabir, Zoe Ostrowski, Shafayet A. Siddiqui, Ian C. Schneider, Long Que

## Abstract

Recent work has highlighted transcranial magnetic stimulation (TMS) as a promising, non-invasive therapeutic approach for a variety of diseases such as Parkinson’s disease (PD). Magnetic stimulation (MS) causes action potentials that result in dopamine release and increases in neuroplasticity. While secretion and the electrophysiology of MS have been well characterized, other downstream effects of MS are unknown. Here we characterize a mechanotransduction pathway that is activated in response to MS in a dopaminergic cell line (N27-A). Integrin tension was measured using DNA-based integrin tension sensors (ITSs), and cytoskeletal and focal adhesion structure was quantified in response to pharmacological inhibitors and activators of common mechanotransduction pathways. MS enhances ITS rupture in N27-A cells in a dose dependent manner. Additionally, both pulse frequency and number control ITS rupture. The ITS rupture that MS causes occurs through both cell migration-independent and -dependent mechanisms. Furthermore, MS enhances integrin forces in N27-A cells through a myosin II- and FAK-dependent pathway by creating more internal adhesions with increased phosphorylated paxillin. This work definitively demonstrates that MS can cause robust changes in mechanotransduction, which likely regulate neuroplasticity and provide an additional mechanism to explain the efficacy and potential of TMS therapy.

TOC graphic

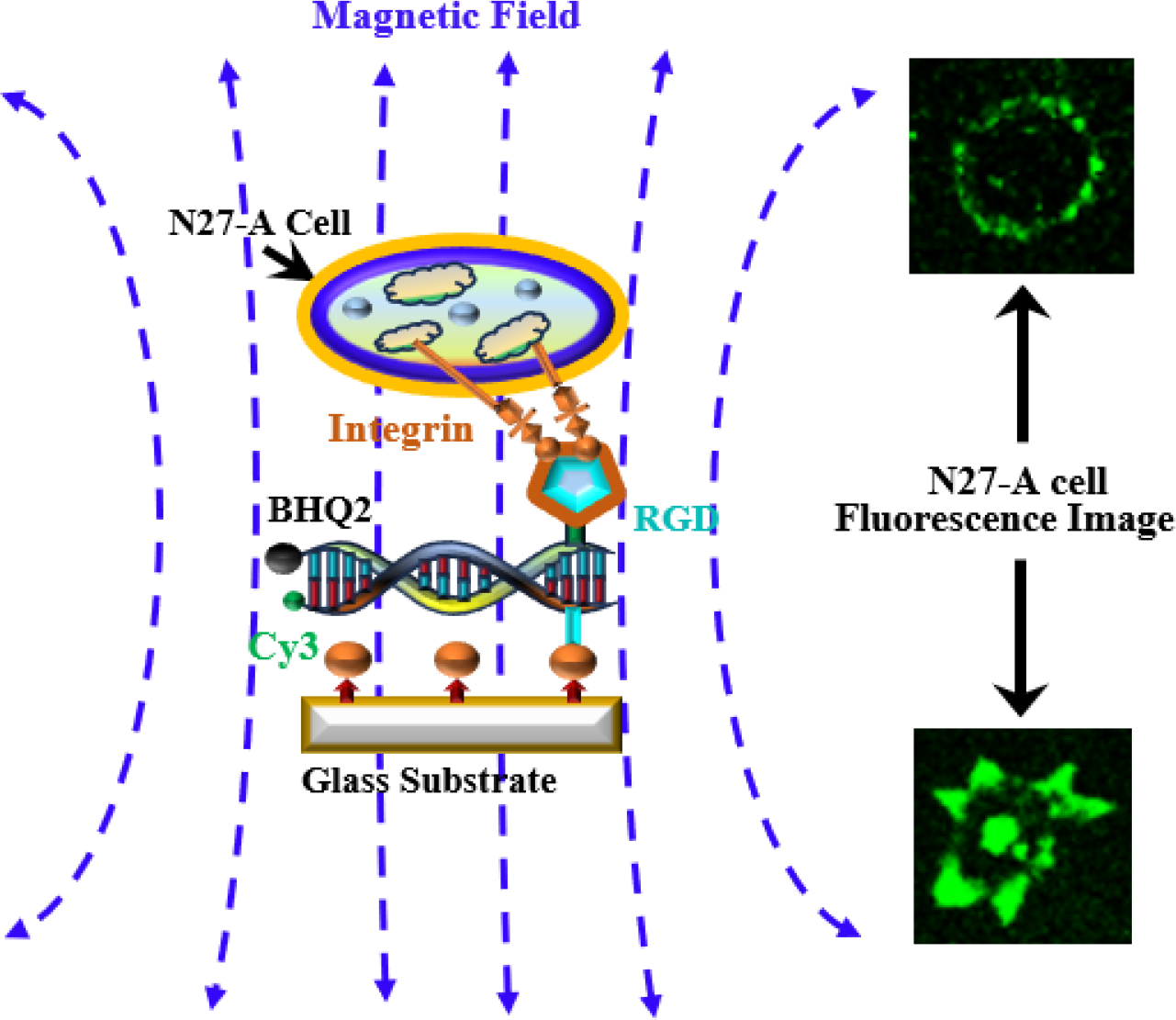

## 1. Introduction

Transcranial magnetic stimulation (TMS) has been approved to treat a variety of diseases such as major depressive disorder (MDD), obsessive compulsive disorder (OCD), migraines and smoking addiction. There is also strong evidence that it could be used to treat anxiety, bipolar disorder and post-traumatic stress disorder (PTSD) as well as Parkinson’s disease (PD), Alzheimer’s disease (AD), chronic pain and fibromyalgia^1–3^. TMS results in the secretion of dopamine and magnetic stimulation (MS) *in vitro* causes growth factor secretion like glial-derived neurotrophic factor (GDNF)^4–6^. Dopamine release induces neuroplasticity^7^, whereas GDNF indirectly supports dopamine release by encouraging dopaminergic neuron survival^8^. The upstream mechanism driving the therapeutic effects includes the stimulation of action potentials in the axons of neurons as well as the release of neurotransmitters, inducing neuroplasticity^9^. Neuroplasticity is driven by synaptic or dendritic plasticity (altering of the structure of the neuronal connections) and neurogenesis (growth of new neurons) as well as functional plasticity (the changing of structures to be used by alternative brain functions). Synaptic or dendritic plasticity and neurogenesis are governed in part by the secretion of factors such as GDNF, but also by mechanobiology, whereby neuronal cells sense and transmit forces to the extracellular matrix (ECM). This results in axonal extension, dendritic spine formation and maturation and neuronal cell migration^10–11^. While it is known that mechanobiology regulates neuroplasticity, nothing is known about how TMS might alter mechanobiological responses to control neuroplasticity.

The induction of dopamine as well as the increase in neuroplasticity have motivated the exploration of TMS as a PD therapy^2^. TMS offers promising, non-invasive symptom relief in PD, particularly for motor symptoms and depression^2, 12–14^. A subclone of the N27 rat dopaminergic neural cell line (N27-A cells) along with its parental line, is commonly used as an experimental model to investigate the mechanisms underlying PD due to its dopaminergic characteristics^15–16^. N27 cells are derived from mesencephalic tissue (midbrain) rat embryo. These cells do not have axons or dendrites and as such likely represent a post-mitotic, but not fully differentiated neuron that has been immortalized. N27 cells seem to express at least some neuronal markers. One study indicates that parental N27 cells do not express active calcium channels for depolarization^17^, but other work indicates the presence of both sodium and calcium channels^18^ and N27-A cells readily depolarize in response to potassium^15^, indicating these cells retain many of the electrophysiological markers of neurons. In previous work, we have explored the effects of MS on N27 dopaminergic neuronal cells^6,19^. The results indicate that MS promotes the proliferation of these cells, with increased expression of GDNF. While MS induces proliferation and survival through secretion of factors in N27 cells, other effects of MS such as those regulating mechanobiology are not known.

Mechanobiology is the field of study concerned with how cells transmit and sense forces and is linked to the ability of the cell to sense the mechanical properties of the surrounding ECM. Forces are sensed and transmitted to the ECM by physical contact of integrins to the ECM^20–22^. These forces can be measured by DNA-based integrin tension sensors (ITSs)^23–25^ or traction force microscopy^26^. Integrins cluster and form focal adhesions, recruiting Focal Adhesion Kinase (FAK) and paxillin. Focal adhesion assembly, maturation and turnover are modulated through myosin II-mediated contractility on the F-actin cytoskeleton^27^. Integrin force transmission to the ECM, mediated by focal adhesions and the cytoskeleton, results in mechanical signaling that affects a broad array of cell functions such as proliferation, differentiation and cell migration. In neurons, these same mechanobiological pathways are needed for dendritic spine assembly and maturation, constituting the link between mechanobiology and neuroplasticity^28–31^. There are a few studies showing that changes in ECM stiffness alter differentiation of stem cells into dopaminergic neurons as well as survival of dopaminergic neurons^32–34^, but this is the extent of the exploration of mechanobiology in dopaminergic neurons. In other neurons, traction forces generated by integrins or other surface adhesion receptors are important in both axonal growth cone and neuronal migration^35–36^. However, no work has explored how MS alters mechanobiology in dopaminergic neuronal cells.

In this work, we characterize the response of N27-A cells, a dopaminergic cell-line model of PD, to MS at the level of mechanobiological outputs. We measure changes in integrin tension sensor rupture in response to MS and show dependence on MS dose, frequency and pulse number. MS enhances integrin tension sensor rupture in both cell migration-dependent and -independent manner. Furthermore, we have identified the myosin II-FAK-paxillin signaling axis to be the driver of increases in integrin tension sensor rupture in response to MS. This same pathway is used in dendritic spine assembly and maturation and demonstrate a potential explanation for the enhanced neuroplasticity observed in response to MS. The characterization of this mechanobiological response to MS supports TMS treatment in other diseases where neuroplasticity is desired or who have specific deficiencies in mechanotransduction as the cause of the disease.

## 2. Materials and Methods

### 2.1. Chemicals and materials

#### N27-A Cell Culture

The N27-A dopaminergic neuron-derived cell line from rat was purchased from Millipore (Cat. No. SCC196). Thawed cells were cultured in RPMI-1640 medium (Sigma Cat. No. 11875-093) supplemented with 10% fetal bovine serum (Gibco Cat. No. 10438-026) and 100 U/ml penicillin and streptomycin (Gibco cat. No. 15140-122). Cells were allowed to grow in the incubator at a regular culture condition of 37 °C with 5% CO_2_ to 80-90% confluency on the cell culture dish and never passed beyond passage 12.

#### Integrin tension sensors (ITSs)

Phosphate Buffered Saline (PBS) was purchased from Thermo Fisher Scientific (Waltham, MA, USA). Glass bottom petri dishes (Product #: D35-10-0-N) were obtained from Cellvis (Sunnyvale, CA, USA). BSA-biotin (Catalog #: A8549) was purchased from Sigma-Aldrich, Fibronectin (Catalog #: 1918-FN-02M) was purchased from R&D Systems, Inc., and Neutravidin (Catalog #: 31000) was purchased from Thermo Fisher Scientific.

### 2.2 Operational principle of ITSs

The working principle of an integrin tension sensor (ITS) is shown in **Figure 1(a)**. The glass substrate is sequentially coated with BSA-biotin mixed with fibronectin, followed by neutravidin and ITS-biotin. A double-stranded DNA (dsDNA) fragment is immobilized via the lower strand, while an RGD peptide, attached to the upper strand, serves as a ligand for integrin receptors on N27-A cells. Integrins bind to the RGD motif, generating traction forces during cell attachment and contraction. At low or no tension, the dsDNA remains intact, keeping the Cy3 fluorophore quenched by the nearby BHQ2, resulting in minimal fluorescence. When the integrin-generated force exceeds the threshold, the dsDNA separates, releasing Cy3 from the quencher and activating fluorescence. Once activated, the ITS maintains its fluorescent signal, enabling real-time imaging of integrin-mediated tension in N27-A cells.

**Figure 1.**
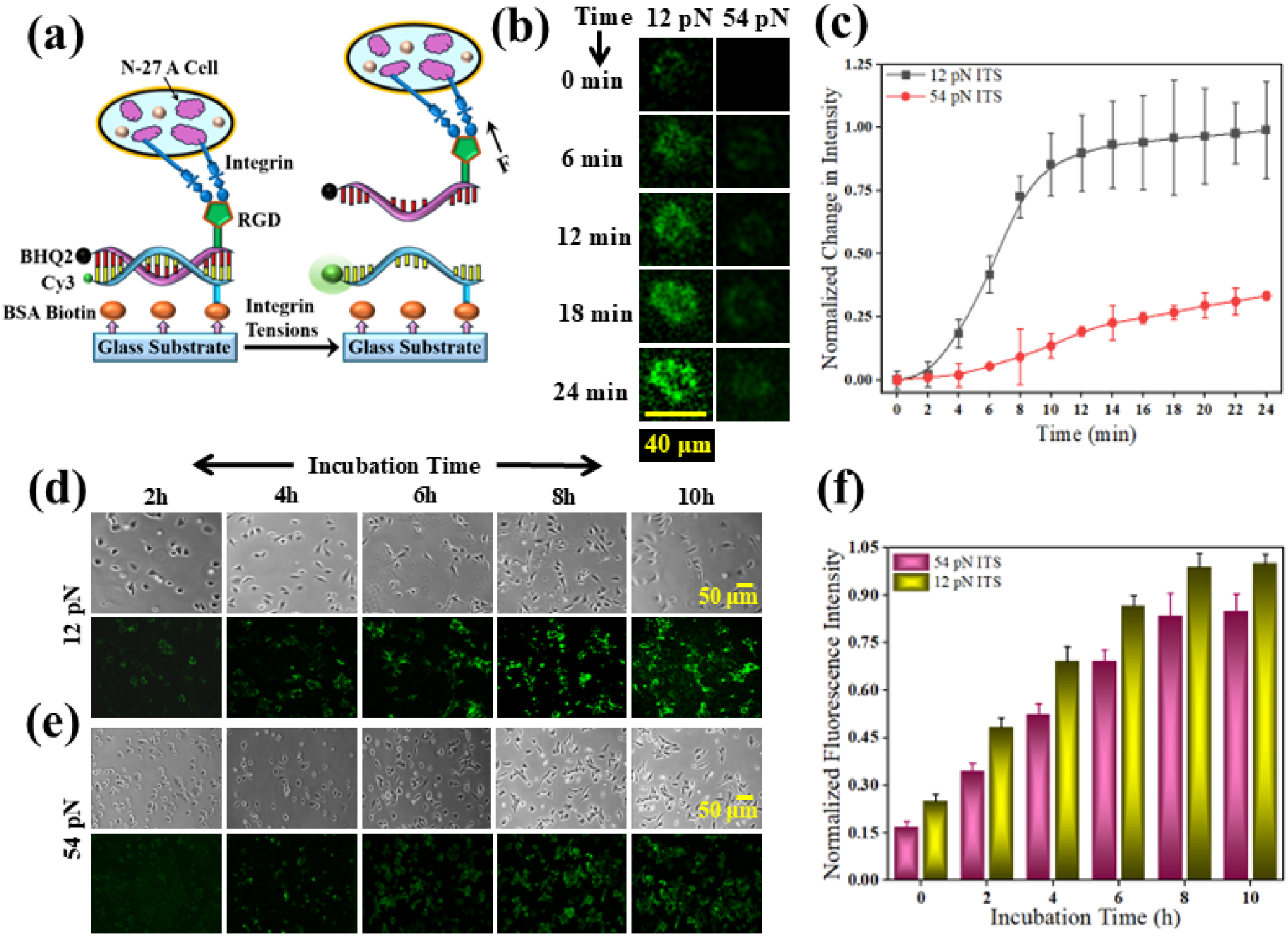
(a) Schematics showing the integrin tension sensor (ITS) structure and its operating principle by binding with a N27-A cell through integrins; (b) Representative fluorescence images on one individual N27-A cell during spreading with 12 pN ITS and 54 pN ITS; (c) Normalized fluorescence intensity changes with incubation time with 12 pN ITS and 54 pN ITS; (d) Representative phase contrast images of N27-A cells plated on 12 pN ITS (top row) and the corresponding fluorescence image (bottom row); (e) Representative phase contrast images of N27-A cells plated on 54 pN ITSs (top row) and the corresponding fluorescence image (bottom row); (f) Normalized fluorescence intensity of 12 pN and 54 pN ITS. Error bars are standard deviation values. For single cell experiments (c): *N_days_* = 3, *N_total experiments_* = 3, *N*_cells per experiment_ = 10. For multi-cell experiments (f): *N_days_* = 3, *N_total experiments_* = 3, *N*_images per experiment_ = 3.

### 2.3 Immobilization of ITSs on coverslip

#### Integrin tension sensor (ITS) design

The tension sensor was engineered using double-stranded DNA (dsDNA). One strand was modified with a fluorophore and biotin, while the complementary strand was labeled with a quencher and an RGDfK ligand. For additional technical details, please refer to our previous publication^24^. The DNA sequence used in the sensor is provided below.

ITS with 12 pN threshold (with Cy3-BHQ2 pair):

5‘-/RGD/GGG CGG CGA CCT CAG CAT/BHQ2/ −3’

5‘-/Cy3/ATG CTG AGG TCG CCG CCC/Bio/ −3’

ITS with 54 pN threshold (with Cy5-BHQ2 pair):

5‘-/RGD/GGG CGG CGA CCT CAG CAT/BHQ2/ −3’

5‘-G/Cy5/T/Bios/ATGATGAGGTCGCCGCCC/−3’

#### ITS immobilization to the coverslip surface

The glass surface was first coated with 100 μg/mL BSA-biotin for 30 minutes, followed by incubation with 100 μg/mL neutravidin for another 30 minutes. Subsequently, the biotin-tagged integrin tension sensor (ITS) was introduced at a concentration of 0.1 μM. Phosphate-buffered saline (PBS) washes were performed between each step to remove unbound components.

### 2.4 Seeding of N27-A cells on the ITSs-coated coverslips

#### Seeding cells

N27-A cells were trypsinized with 0.05% trypsin/EDTA and counted. Then 80,000 cells were incubated with 2 mL media on one side of 22 × 22 mm glass coverslip (Corning Cat. No. 2850-22) surface of previously modified with attached 12 pN or 54 pN ITSs, respectively.

#### Making cell chambers

It takes around 4 hours for cells to attach on the glass surface and begin their spreading phase. After 4 hours of incubation, a cell migration chamber was made and filled with clear RPMI media (ThermoFisher Cat. No. 11835030) supplemented with 10% fetal bovine serum, 1% penicillin and streptomycin, and 15 mM 4-(2-Hydroxyethyl)piperazine-1-ethanesulfonic acid (HEPES). The cell migration chamber was made from a one-sided sticky soft silicone rubber sheet (Amazon, Laimeisi brand) where approximately 22 × 22 mm size of silicone rubber sheet was cut in Cricut with the internal hollow chamber size of 20 × 20 mm and the prepared chamber was attached on a 22 × 40 mm glass coverslip (Corning Cat. No. 2975-224). During N27-A cell migration the chamber was filled with warm (37 °C) clear RPMI media, and the glass slide of modified ITS surface attached with N27-A cells was placed on the chamber facing down so that the cells were in contact with clear RPMI media then the chamber was sealed with melted VALAP. Live cell migration was recorded in the microscope for 8 hours at 5-minute intervals.

### 2.5 Stimulating N27-A cells with MS

#### Control experiments

In these experiments, either MS or inhibitor was not applied to the N27-A cells after the glass substrate was prepared with BSA-biotin mixed with fibronectin, neutravidin, and ITS-biotin. The cells were incubated overnight at 37 °C in the incubation chamber, and fluorescence intensity was measured the following day without any MS treatment.

#### MS experiments

After the glass surface was prepared with BSA-biotin mixed with fibronectin, neutravidin, and ITS-biotin, N27-A cells were introduced into it followed by overnight incubation at 37 °C in a chamber where CO_2_ is available. After incubating it overnight, the cells were stimulated with the MS device.

#### Inhibitor Experiments

The glass surface was prepared with ITSs as described above. N27-A cells were seeded and given 1 hour to attach and spread. After 1 hour, the cells were treated with either DMSO (control), 10 µM defactinib (Tocris Bioscience 7305), 10 µM blebbistatin (Sigma) or 10 µM omecamtiv mecarbil (Selleckchem S2623). Either no MS treatment or one MS treatment at 73 mTs immediately followed the addition of the inhibitor. Images were taken at 20 x (*NA* = 0.5, Nikon). every hour for 4 hours.

### 2.6 Immunofluorescence Imaging and Analysis

#### Staining Cells

Glass coverslips were coated with 100 µl of 10 µg/ml fibronectin for 30 minutes. Then 100,000 N27-A cells were seeded onto the coverslips and allowed to attach and spread for 1 hour. Then the cells were treated with either DMSO (control), 10 µM defactinib, 10 µM blebbistatin, or 10 µM omecamtiv mecarbil. Either no MS treatment or one MS treatment at 73 mTs was done immediately following the addition of inhibitor. Four hours after the inhibitor was added the cells were fixed for 10 minutes at 37 °C using 4% paraformaldehyde (Fisher Scientific) prepared by diluting 16% paraformaldehyde in cytoskeleton buffer. The cytoskeleton buffer was composed of 10 mM 2-N-morpholino ethane sulfonic acid (MES buffer pH 6.1) (Fisher Scientific), 3 mM MgCl_2_ (Fisher Scientific), 138 mM KCl (Fisher Scientific), and 2 mM EGTA (Sigma Aldrich). Cells were then treated with 100 mM glycine (Fisher Scientific) for 15 minutes. Cells were washed 3 times with 1X Tris Buffered Saline (TBS) for 5 minutes each. Cells were first stained with Alexa 488-phalloidin (Thermo Fisher Scientific) for 1 hour in 1x TBS containing 0.1% (v/v) Tween-20 (Fisher Bioreagents) and 2% (w/v) bovine serum albumin (BSA, Sigma Life Sciences). After 1X TBS washing, cells were treated with 1:200 dilution of anti-paxillin (BD Transduction, mouse, 610052) and 1:100 dilution of pY118 paxillin primary antibody (Invitrogen rabbit, 44722G) for 1 hour. Samples were washed with 1X TBS three times for 5 minutes each. Next, the samples were treated with 1:400 dilution of donkey anti-rabbit 568 secondary antibody (Jackson ImmunoResearch), 1:400 dilution of donkey anti-mouse Cy5 secondary antibody (Jackson ImmunoResearch) and 1:800 dilution of DAPI for 1 hour. Samples were imaged using a 100x objective (*NA* = 1.49, Nikon).

#### Imaging Stained Cells

Images were taken on a Moment CMOS (Teledyne) by focusing on the bottom of the cell. Images were flattened by first subtracting the offset of the microscope (∼100 gray levels). Then the image was divided by a reference image of soluble fluorescent material and multiplied by the maximum fluorescent value of the reference image. Individual cells were manually outlined to create a total cell area. The outline was uniformly shrunk between 1.3 and 3.3 µm to exclude the larger external focal adhesions on the edge of the cell creating an internal cell area, and the external area was the area excluded from the internal cell area. The paxillin images were background subtracted using a rolling ball background subtraction (10 pixels) and then the threshold was adjusted to see the focal adhesions and saved as regions of interest (ROIs). These ROIs were used to create a focal adhesion mask for each cell. The internal, external and total areas were overlaid onto the mask to make an internal, external and total focal adhesion mask. These masks were used to measure the total focal adhesion area per cell and the integrated fluorescence of paxillin and pY118 paxillin per cell.

### 2.7 Data collection and analysis

An epifluorescence microscope (Nikon TiE) equipped with Cy3-specific filters and a Moment CMOS (Teledyne) was used to capture fluorescence images of the N27-A cells. The fluorescence intensity of cells without TMS treatment was measured and used as a baseline reference, while the cells exposed to TMS served as the positive control group. After imaging, the collected data was processed and analyzed using ImageJ software. For statistical reliability, fluorescence intensity values were averaged from at least three independent experiments to ensure consistency and reproducibility of the results. Normalized fluorescence intensity = (*I* - *I_b_*)/(*I_max_* - *I_b_*), where *I* is the mean fluorescence intensity, *I_max_* is the maximum fluorescence intensity and *I_b_* is the mean background fluorescence intensity. For migration measurements, normalized change in cell migration area = (*A* - *A*_0_)/(*A_max_* - *A*_0_), where *A* is the mean cell migration area after a particular incubation time, *A****_0_*** is the mean cell migration area at the initial condition and *A_max_* is the maximum cell migration area.

## 3. Results and Discussion

### 3.1 ITS rupture dynamics in N27-A cells over short and long timescales

As shown in **Fig. 1(a)**, when N27-A cells adhere to the surface coated with ITS, integrins on the cell membrane bind to the DNA-based tension sensors through the RGD sequence. As an integrin exerts mechanical force, rupture of the double-stranded DNA (dsDNA) structure within the ITS leads to the separation of the fluorophore and quencher pair attached to the DNA strand, thereby producing a detectable fluorescence signal. This fluorescence emission serves as an indicator of molecular tension exerted at a loading rate over time generated by integrin-ligand interactions at the cell– substrate interface.

To monitor the dynamic process of integrin force generation, fluorescence images were taken for individual N27-A cells after being plated onto ITS-functionalized surfaces and are shown in **Fig. 1(b)**. The timing of these images spans the initial spreading time of these cells. ITS fluorescence was quantified, normalized and shown in **Fig. 1(c)**. The fluorescence signal intensities gradually increase over time, reflecting a time-dependent rise in the number of ITS ruptured by integrin force transmission. Notably, the fluorescence signal recorded from ITS with a tension threshold of 12 pN is significantly higher than that from the 54 pN ITS, indicating that the majority of integrin forces exerted by N27-A cells are only sufficient to rupture the lower-threshold 12 pN ITS but not the higher-threshold 54 pN ITS. Eventually, the fluorescence intensities reach a plateau, typically within 10 to 20 minutes after cell seeding. This plateau phase implies that cell spreading has ceased and the majority of integrins on the cell surface have either formed stable bonds or that the available sensors have been maximally ruptured.

To investigate the dynamics of integrin force generation post-spreading, fluorescence images were taken at longer timescales (2-10 h) in a sampling of regions for both 12 pN and 54 pN ITSs and are shown in **Figure 1(d-e)**. In contrast to the rapid saturation observed during spreading, the fluorescence intensity increases gradually up to 8 hours of incubation in **Figure 1(f)**. This sustained fluorescence increase followed by another steady state at around 8 hours shows fluorescence levels of 54 pN and 12 pN ITS sensors that are more evenly matched. Migration that occurs after spreading likely results in cells rupturing ITS in new regions as the cells migrate, causing the increase in fluorescence. These findings show that early during spreading low forces (12 pN) are primarily exerted and these forces come to a pseudo-steady state at around 15 minutes. As cells begin to migrate, the forces of 12 pN and 54 pN are more evenly balanced, increasing over time as cells move over the ITS substrate.

### 3.2 TMS enhances ITS rupture in N27-A cells

Having established the ability to detect ITS rupture under N27-A cells during spreading and migration, we were interested in determining whether MS produces changes in ITS rupture. The schematic diagram of the MS experimental setup, along with the timeline for MS application on N27-A cells, is presented in **Figure 2(a)** and **Figure 2(b)**, respectively. In all subsequent experiments, MS was administered after the N27-A cells had been seeded onto coverslips and incubated overnight. The overnight incubation ensured sufficient time for initial cell adhesion before the application of magnetic stimulation.

**Figure 2.**
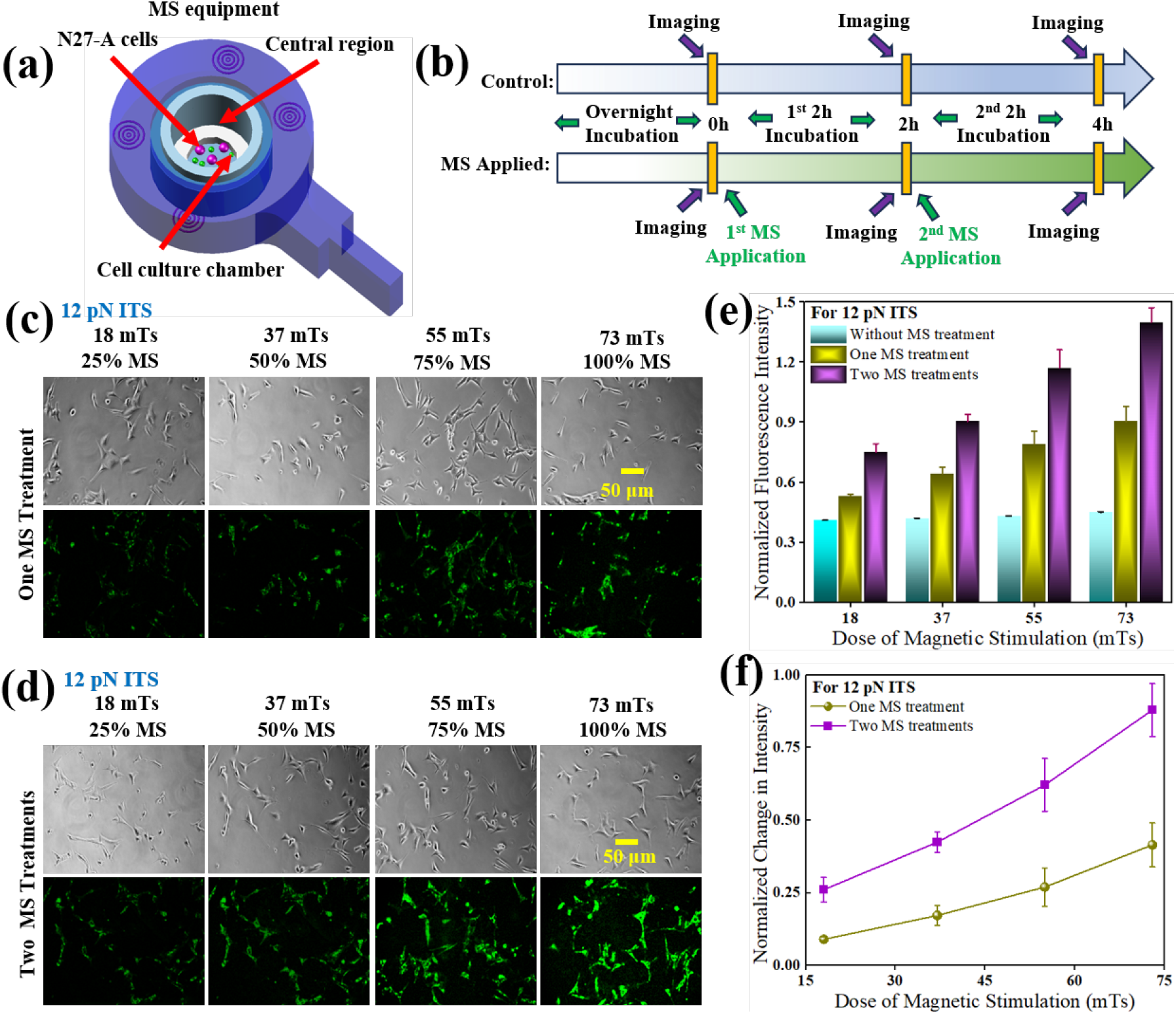
(a) Sketch showing the MS treatment on N27-A cells cultured in a tissue culture dish; (b) Time line identifying the MS treatments and fluorescence images taken from experimental samples and control samples; (c) Representative phase contrast images of N27-A cells with 12 pN ITS (top row) and the corresponding fluorescence images (bottom row) subjected to one MS treatment of varied strength; (d) Representative phase contrast images of N27-A cells with 12 pN ITS (top row) and the corresponding fluorescence image (bottom row) subjected to two MS treatments of varied strength; (e) Normalized fluorescence intensity of the control sample and the samples subjected to one MS treatment and two MS treatments; (f) Net changes in fluorescence intensity after subtraction of the control sample values. Error bars are standard deviation values. *N_days_* = 3, *N_total experiments_* = 3, *N*_images per experiment_ = 3.

To evaluate the effects of different MS doses on integrin force, a series of experiments was conducted using varying doses of magnetic stimulation. The measured magnetic field strength at the center of the ring-shaped MS device (**Figure 2(a)**) is 730 µT, representing 100% of its output^6^. The magnetic dose is calculated using the formula: Magnetic dose = magnetic field strength (T) x exposure time (s)^37^. In these experiments, each MS session delivers 100 magnetic pulses at a magnetic pulse repetition frequency (mPRF) of 0.1 Hz. Representative phase contrast images and fluorescence images of N27-A cells plated on 12 pN ITS for these conditions are shown in **Figures 2(c)** and **2(d)**, corresponding to one-time and two-time MS treatments, respectively. In both cases, the fluorescence signals, indicative of integrin force transmission, show a clear dose-dependent increase with higher doses of MS (**Figure 2(e-f)**). Furthermore, two MS treatments yield significantly higher fluorescence signals than a single MS treatment. Much smaller responses were observed using the 54 pN ITS (**Supplemental Figure 1**), suggesting that MS activates lower tension integrin responses. Collectively, these findings demonstrate that the magnitude and number of MS treatments can modulate the integrin force exerted by N27-A cells and that TMS primarily upregulates integrin tension that is at or below 54 pN.

To determine if MS initiates a process that results in progressive increases in fluorescence significantly past the time of stimulation, different numbers of MS treatments were applied to N27-A cells. Each MS session delivers 100 magnetic pulses at a mPRF of 0.1 Hz. Following each MS treatment, the cells were incubated for either one or two hours, after which fluorescence imaging was conducted to assess the level of integrin force transmission through fluorescent signal intensity. **Figure 3(a)** presents the fluorescence images of the control samples, which did not receive any MS treatment and therefore serve as a baseline for comparison. **Figure 3(b)** shows the fluorescence images taken after a single MS treatment, while **Figure 3(c)** illustrates the results obtained after one to four successive MS treatments. The fluorescence signal becomes progressively more intense with an increasing number of MS treatments. The fluorescence intensities were quantified to provide a clearer demonstration of this observation. In **Figure 3(d)**, the quantified fluorescence intensities are plotted for N27-A cells that received three TMS treatments, each with different MS doses. This analysis helps to distinguish the effects of not only the number of treatments but also the dose applied during the stimulation process. Furthermore, **Figure 3(e)** provides a comparative overview of the quantified fluorescence intensities across different experimental conditions, namely, control samples, samples that received a single MS treatment, and those that underwent four treatments. Interestingly, fluorescence does not increase past about 1 hour after stimulation. Overall, the results strongly support that MS dose regulates integrin forces, producing increases in ITS rupture for larger doses. This response is more dramatically observed using the 12 pN ITS in comparison to the 54 pN ITS. Furthermore, increases in integrin forces occur soon after stimulation, but wane after stimulation.

**Figure 3.**
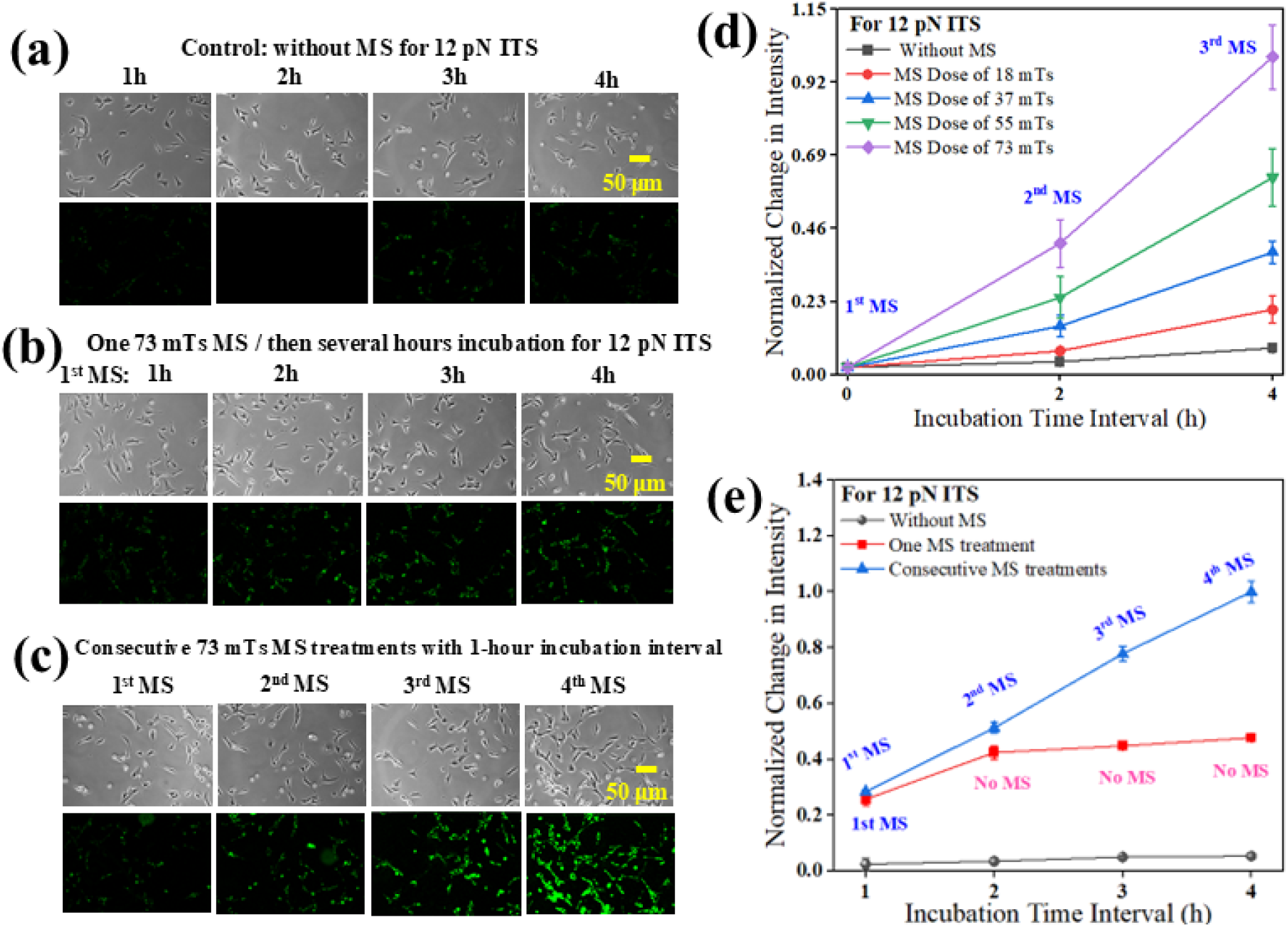
Representative phase contrast images of N27-A cells with 12 pN ITS and the corresponding fluorescence images subjected to MS treatments; (a) Control samples without any MS treatments; (b) One MS treatment followed by 4 hour incubation; (c) Four consecutive MS treatments with 1-hour incubation interval; (d) Normalized fluorescence intensity of the samples with 12 pN ITS subjected to consecutive MS treatments of varied MS strength with a 1 hour incubation interval; (e) Normalized fluorescence intensity of the samples with 12 pN ITS with different numbers of MS treatments with a 1 hour incubation interval. Error bars are standard deviation values. *N_days_* = 3, *N_total experiments_* = 3, *N*_images per experiment_ = 3

### 3.3 TMS pulse frequency and number controls regulate ITS rupture in N27-A cells

Having established that MS increases ITS signal by enhancing integrin force in a MS dose-dependent manner, we sought to explore whether other tunable characteristics of MS used in clinical applications such as pulse frequency and number affect integrin forces. Consequently, in this series of experiments, mPRF ranging from 0.05 Hz to 0.2 Hz were used to stimulate N27-A cells. For each treatment condition, 50 TMS pulses were administered. Representative fluorescence images corresponding to the different mPRFs are presented in **Figure 4(a)**, while the quantified fluorescence intensities are shown in **Figure 4(b)**. The ITS signal and consequently integrin force were not linearly dependent on frequency. Below the base case of 0.1 Hz, ITS signal fell off dramatically in comparison to frequencies above 0.1 Hz, which seemed to result in approximately linear increases in ITS signal with frequency. Pulse number (a different way of altering dose) was also explored, so an additional set of experiments was conducted in which the mPRF was held constant at 0.1 Hz, while the number of MS pulses was varied from 25 to 100. Following each treatment, the N27-A cells were incubated for two hours before fluorescence imaging. Representative images captured under these conditions are displayed in **Figure 4(c)**, and the corresponding quantified fluorescence intensities are plotted in **Figure 4(d)**. Similar to the experiments altering dose above, ITS fluorescence intensity increases with a greater number of MS pulses in a linear fashion. Consequently, integrin force increases with increased MS stimulation frequency and this response is linear above 0.1 Hz. Also, linear increases in integrin force occur for a higher number of pulses, which aligns with linear increases seen with more stimulation periods and higher stimulation strength.

**Figure 4.**
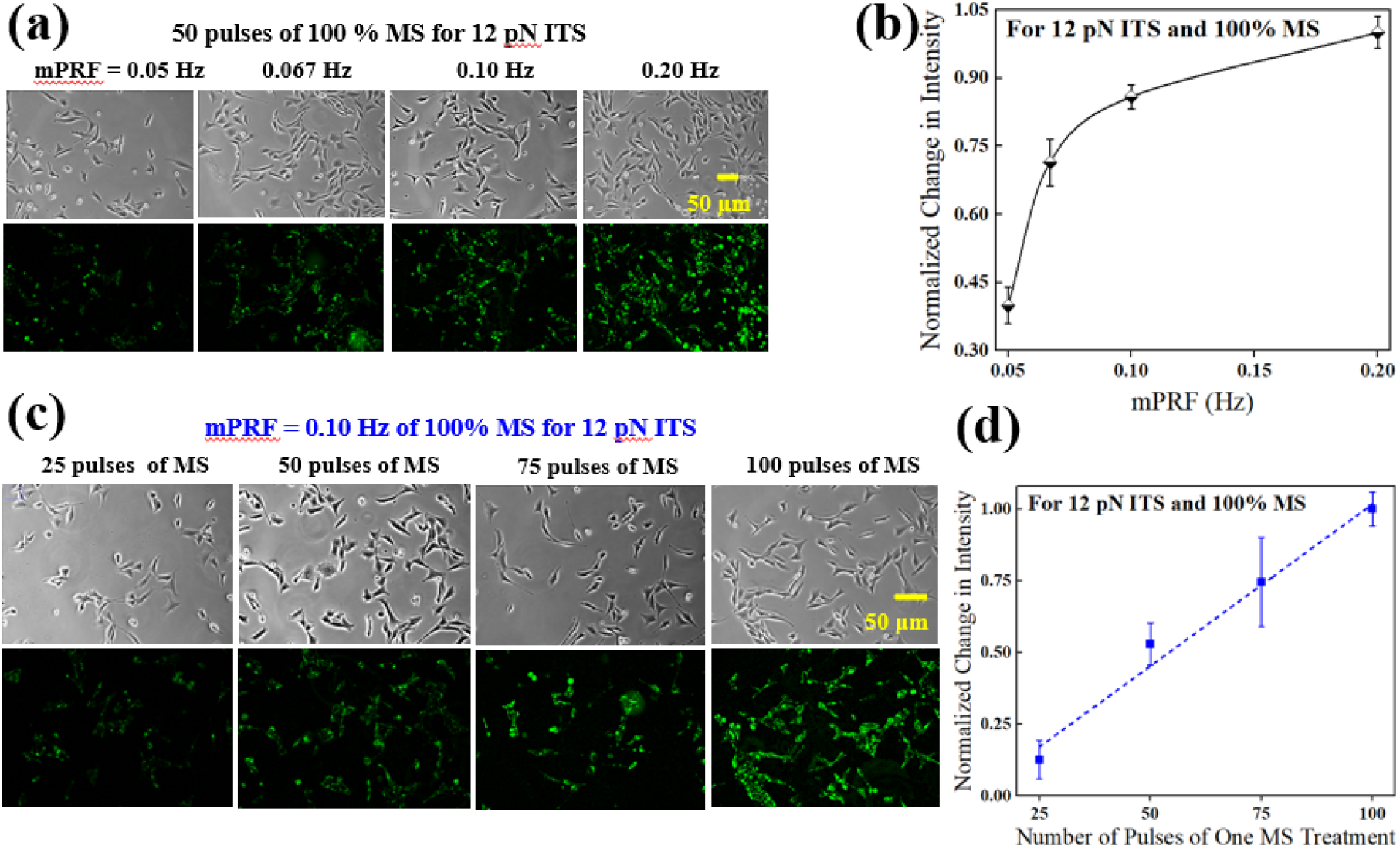
(a) Representative phase contrast images of N27-A cells with 12 pN ITS (top row) and the corresponding fluorescence image (bottom row) subjected to a single MS treatment at a different mPRF, followed by a 2 hour incubation; (b) Normalized fluorescence intensity for (a). (c) Representative phase contrast images of N27-A cells with 12 pN ITS (top row) and the corresponding fluorescence image (bottom row) subjected to a single MS treatment with varying magnetic field pulse numbers, followed by a 2 hour incubation; (d) Normalized fluorescence intensity for (c). Error bars are standard deviation values. *N_days_* = 3, *N_total experiments_* = 3, *N*_images per experiment_ = 3.

### 3.4 MS enhances integrin rupture through both N27-A cell migration-independent and - dependent mechanisms

The ITS fluorescence intensity increases over both short timescales (30 min) and longer timescales (10 h) under N27-A cells, indicating increases during both adhesion, which occurs over minutes initially and migration, which occurs over hours. Additionally, MS stimulation leads to larger ITS signals under N27-A cells and this response is observed to be more robust using the 12 pN ITS. This suggests that MS increases integrin force generation by N27-A cells. However, two different mechanisms could produce this result. First, MS could increase the tension or loading rate at each integrin resulting in more integrins per unit area that are exerting enough force to rupture the ITS and generate a fluorescence signal. This could occur in non-dynamics cells that are not moving (***<u>integrin tension/loading mechanism</u>***). Second, TMS could be increasing the number of integrins per unit time that bind to and rupture ITSs, while not increasing the tension at each integrin. This could occur either during dynamic adhesion or membrane movements or during migration (***<u>integrin dynamics</u> <u>mechanism</u>***). Both scenarios result in what we describe above as higher integrin forces (as opposed to integrin tension, which we use to describe single molecule events) or loading rates. To determine which mechanism the cell is using, we examined ITS signals with and without MS treatment in individual N27-A cells. Treatment was done immediately after plating or well after cells were plated to probe whether increases in integrin force or loading occurred during cell spreading or during cell migration. We also probed ITS fluorescence increases over short (min) and long (h) timescales. We also examined the migration of N27-A cells with and without MS treatments by tracking the cumulative area that the cells have occupied over time in order to directly compare this to the ITS fluorescence signal over time.

N27-A cells were stimulated with MS and ITS signal was measured over short times after stimulation. First, MS was applied immediately after the N27-A cells were seeded onto the ITS-coated coverslips, without any prior incubation. Following the TMS treatment, the fluorescence imaging was performed on live cells over the timescale of minutes to assess integrin force. The resulting fluorescence images of individual cells, along with the quantified fluorescence intensities, are presented in **Figure 5(a-b)**. TMS enhances the ITS signal very quickly during spreading, well before migration. However, this effect was very similar to what is observed if MS stimulation occurs after the cells have been incubated overnight on the substrate. These results are shown in **Figure 5(c)**. This suggests that this increase in integrin forces does not depend on whether the cell is actively spreading or already spread and likely does not require cell migration, which occurs over longer timescales. This suggests that the integrin tension/loading mechanism is contributing to this response. To more clearly determine whether the integrin tension mechanism or integrin dynamics mechanism is driving increases in ITS fluorescence, we conducted real-time monitoring of the migration behavior of individual N27-A cells and measured cumulative area over long timescales. As illustrated in **Figure 5(d)**, during the initial few hours, the extent of cell migration, quantified by measuring the cumulative area over which the cell has migrated, remained relatively constant, suggesting minimal cell migration at the beginning of the cell seeding. However, at around 4 h, significant migration began, resulting in an increase in the cumulative area that the cells occupied over time, even in the absence of MS. Cells stimulated with MS exhibited a significantly greater increase in migration compared to those that were not subjected to MS. The normalized ITS signal was quantified over the same timescale as migration and is shown in **Figure 5(e)**. Fluorescence increases immediately, even over the first 4 h, when migration is not occurring, suggesting that the tension/loading mechanism (stronger individual integrin bonds or higher loading rate) is contributing to increases in ITS fluorescence. However, between 4 and 8 hour, ITS fluorescence continues commensurate with cumulative area, suggesting that the integrin dynamics mechanism (faster turnover of integrins) also contributes. **Figure 5(f)** presents the relationship between normalized fluorescence intensity and the normalized change in cell migration area. If ITSs were ruptured only because migration increased, this curve would be similar to the cumulative area curve in **Figure 5(d)**. However, early on, before migration occurs more numerous integrin rupturing events were taking place. Taken together, these findings support the hypothesis that MS elevates the tension or loading rate of integrins over short times, but over longer times MS also enhances cell migration and the dynamics of integrin-extracellular matrix binding.

**Figure 5.**
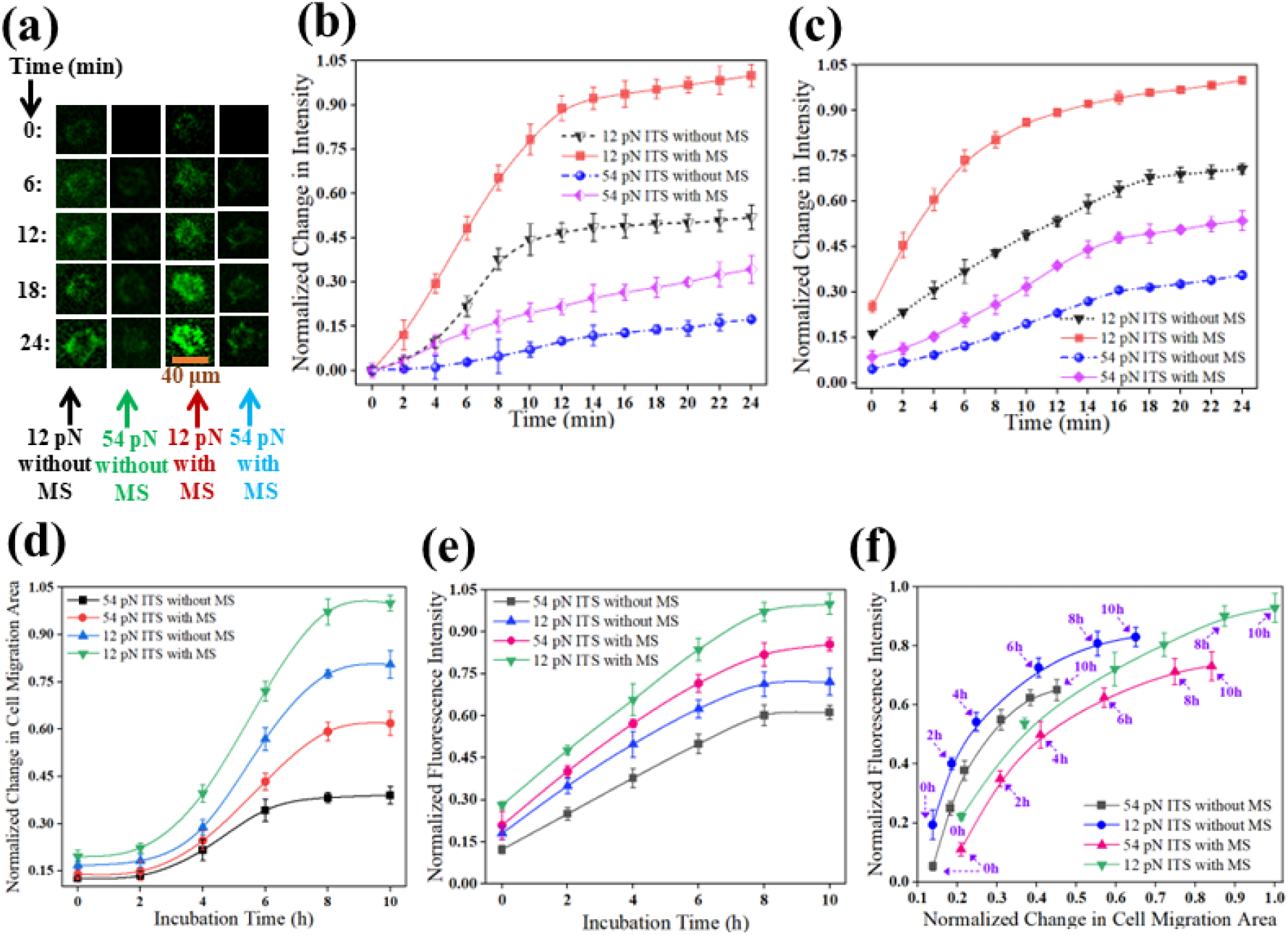
(a) Representative fluorescence images on one individual N27-A cell after overnight incubation on 12 pN ITS and 54 pN ITS with and without MS treatment; (b) Normalized fluorescence intensity changes on individual cells before spreading on 12 pN ITS and 54 pN ITS with and without MS. Time zero is at MS and start of spreading; (c) Normalized fluorescence intensity changes on individual cells after overnight incubation on 12 pN ITS and 54 pN ITS with and without MS. Time zero is at MS and after overnight spreading; (d) Cumulative migration area of N27-A cells during a 10 hour incubation time with and without one MS treatment at time zero. (e) Normalized fluorescence intensity of cells during the same time period as shown in (d). (f) Normalized fluorescence intensity as a function of normalized cell migration area. Error bars are standard deviation values. For single cell experiments (b-d): *N_days_* = 3, *N_total experiments_* = 3, *N*_cells per experiment_ = 10. For multi-cell experiments (e-f): *N_days_* = 3, *N_total experiments_* = 3, *N*_images per experiment_ = 3..

### 3.5 MS enhances integrin forces in N27-A cells through FAK- and myosin-dependent pathways by creating more internal adhesions with increased phosphorylated paxillin

MS enhances integrin forces as observed by ITS fluorescence. Integrins connect the contractile actomyosin directly to the ECM in order to transmit tension that ruptures ITSs. To determine the mechanism behind the increase of integrin forces with MS treatment, three inhibitors were used. N27-A cells plated on 12 pN ITS were treated with 10 μM defactinib, 10 μM blebbistatin, 10 μM omecamtiv mecarbil or DMSO (control) 1 hour after seeding cells. MS treatment immediately followed the addition of the inhibitor. The control cells treated with DMSO showed similar responses to MS as shown in **Figure 6(a)**. The inhibitors showed either enhanced or attenuated ITS fluorescence signal after MS treatment, as shown in **Supplemental Figure 2-5**. Defactinib is a FAK inhibitor. FAK inhibition in nearly all cells disrupts integrin-mediated force transmission through focal adhesions^38–40^. Treatment of N27-A cells with 10 μM defactinib decreases the ITS fluorescence intensity for both the TMS and no TMS treated cells as shown in **Figure 6(b)**. Blebbistatin is a myosin II inhibitor and is known to decrease cell contractility and force generation. Treatment of N27-A cells with 10 μM blebbistatin resulted in a decrease in ITS fluorescence intensity for both MS and no MS treated cells as shown in **Figure 6(c)**. Omecamtiv is a cardiac myosin II activator and is known to increase cell contractility and force generation in cardiac cells ^41^. As shown in **Figure 6(d)**, 10 μM omecamtiv treatment increased the ITS fluorescence intensity for both TMS and no TMS treated cells compared to the control. This was puzzling given these cells do not express cardiac myosin II. While omecamtiv is known to bind other forms of myosin II with low affinity^41^, at higher doses there are some effects on non-cardiac myosin II. In addition, it seems to stimulate voltage-gated Na^+^ channels in neuronal cells^42^. These voltage-gated Na^+^ channels can bind to myosin II^43–44^ and lead to the activation of integrin forces that we observe. To assess which intervention created the largest difference in response to MS under different inhibition conditions, the normalized ITS fluorescence intensity was plotted on the same graph and shown in **Figure 6(e)**. MS treatment appears to increase myosin activity in the cell, resulting in the increased force seen under all inhibitor conditions. Defactinib (*in vitro* FAK activity *IC*_50_ = 0.6 nM) treatment caused both a lower ITS signal as well as a smaller difference (0.080) between MS stimulated and unstimulated conditions than the difference (0.13) seen under blebbistatin (*in vitro* myosin activity *IC*_50_ = 3 μM) treatment. This indicates that FAK is somewhat more important than myosin II in driving the MS increase in integrin tension. However, it is possible that not all myosin II is inhibited. Interestingly, while omecamtiv (cellular Na^+^ channel *IC*_50_ = 2-10 μM) treated cells showed an increase in ITS fluorescence, the difference (0.30) between MS stimulated and unstimulated was smaller than the difference (0.44) in the control, indicating that the ITS signal is beginning to saturate. MS stimulation has been shown to activate voltage-gated Na^+^ channels^45–47^. Consequently, MS and omecamtiv may both induce voltage-gated Na^+^ channels, which has the effect of activating myosin II and FAK downstream to upregulate integrin tension and dynamics within N27-A cells.

**Figure 6.**
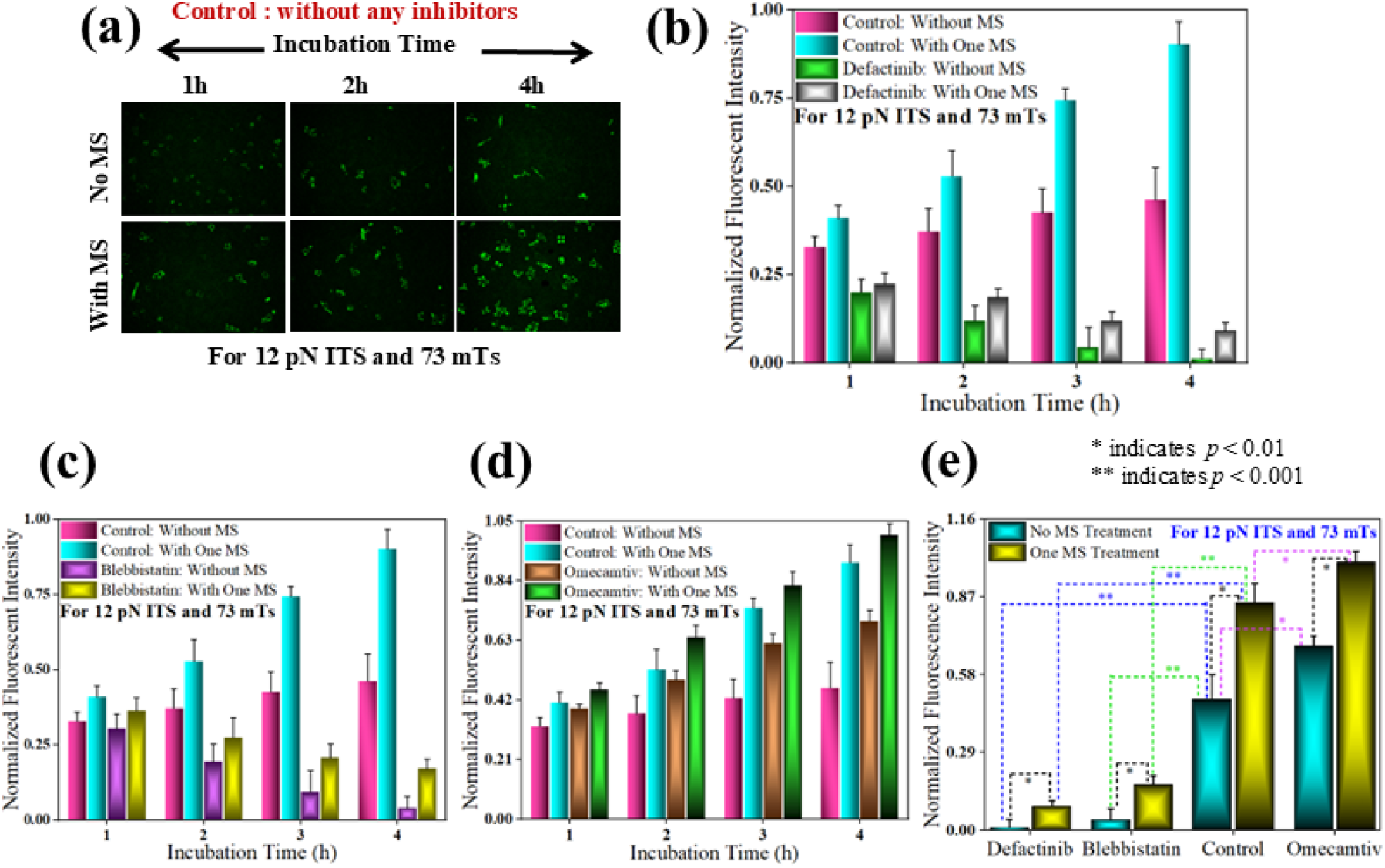
(a) Representative fluorescence images of N27-A cells with 12 pN ITS in the absence of inhibitors, with or without MS treatment of 73 mT at time zero; (b) Normalized fluorescence intensity of N27-A cells with 12 pN ITS with or without 10 μM defactinib, and with or without a single MS treatment of 73 mTs; (c) Normalized fluorescence intensity of N27-A cells with 12 pN ITS with or without 10 μM blebbistatin, and with or without a single MS treatment of 73 mTs; (d) Normalized fluorescence intensity of N27-A cells with 12 pN ITS with or without 10 μM omecamtiv, and with or without a single MS treatment of 73 mTs; (e) Comparison of the normalized fluorescence intensity of N27-A cells between control (without any inhibitors), 10 μM defactinib, 10 μM blebbistatin, and 10 μM omecamtiv with 12 pN ITS and 73 mTs MS. Error bars are standard deviation values. *N_days_* = 3, *N_total experiments_* = 3, *N*_images per experiment_ = 3

To further investigate the hypothesis that MS treatment causes an increase in myosin II activity that results in changes in focal adhesion signaling, N27-A cells were fixed and stained for focal adhesions using antibodies for paxillin and phosphorylated paxillin (pY118). FAK phosphorylates paxillin on Y118, which regulates focal adhesion assembly and turnover^46^. Phosphorylated paxillin on Y118 increases with myosin II activity^48^, while dephosphorylation occurs commensurate with a lowering of myosin II activity and traction forces^49^. Finally, focal adhesion force fluctuations, which drive durotaxis (directed migration in response to stiffness gradients) are dependent on paxillin phosphorylation on Y118^38, 50^. N27-A cells were seeded on fibronectin-coated coverslips, the same ECM molecule used with the ITS in the above studies. Cells were given 1 hour to attach and spread before being treated with MS. Cells were fixed 4 hour after MS treatment and then stained for F-actin, paxillin, pY118 paxillin and DNA in the nucleus. Shown in **Figure 7(a)** and **Supplemental Figure 6**, unstimulated cells had large paxillin-containing focal adhesions at the edge of the cell with few to no focal adhesions directly under the cell. However, MS treated cells had large paxillin-containing focal adhesions near the edge of the cell as well as smaller focal adhesions directly under the cell. Consequently, the cell was manually divided into two regions separated by the red line in the paxillin images shown in **Figure 7(a)**. MS resulted in higher total focal adhesion area per cell, integrated paxillin fluorescence and integrated pY paxillin fluorescence as measured over the entire cell **Figure 7(b-d)**. All of these measures are indicative of cells exerting larger forces and support the above observation that ITS signal and integrin forces are increased. Interestingly, MS produced more robust changes in these focal adhesion measurements when the measurement was constrained to the internal region and more subtle changes were observed in the external region. This seems to be somewhat different than what was reported previously by our group in response to MS, but here we focused solely on single cells so as not to complicate the interpretation with cell-cell contacts^19^ and rather than just measuring focal adhesion number, we quantified the sum area and sum fluorescence, which are better measures of how much adhesion and consequently, integrin-mediated force cells are generating. External adhesions observed in unstimulated cells are associated with static non-motile cells generating forces at the edge, whereas internal adhesions observed in MS cells are associated with dynamic, migrating cells. Consequently, MS results in more focal adhesions that are driven primarily by an increase in internal focal adhesions that contain more phosphorylated paxillin on Y118. These focal adhesion characteristics are indicative of fast moving cells, supporting the integrin dynamics mechanism of increased integrin force as measured by increased ITS fluorescence after MS.

**Figure 7.**
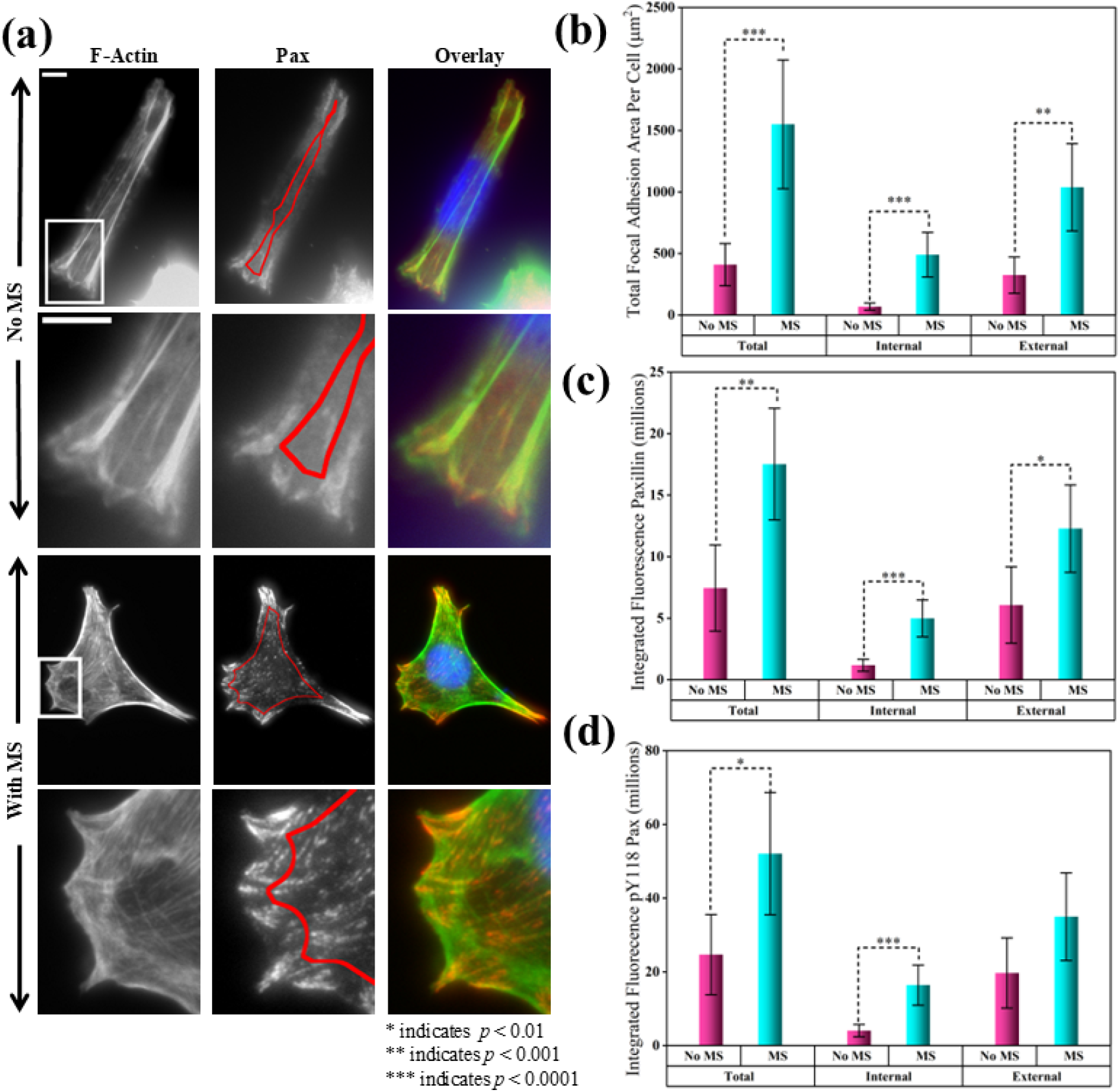
(a) Fluorescence images of N27-A cells on fibronectin coated coverslips stained for F-actin and paxillin with and without MS treatment. Overlays of F-actin, paxillin and DAPI are included. Calibration bar length = 5 μm; (b) Total focal adhesion area per cell in the internal, external and total cell areas designated by the red line on the paxillin image in (a); (c) Integrated fluorescence per cell of paxillin in focal adhesions in the internal, external and total cell regions; (d) Integrated fluorescence per cell of pY118 paxillin in focal adhesions in the internal, external and total cell regions. *N_days_* = 2, *N_total experiments_* = 2 and *N_cells per experiment_* = 10. Error bars are 95% confidence intervals. Asterisks * and ** and *** indicate statistically significant differences with *p* < 0.01, 0.001 and 0.0001 respectively using two-tailed student’s *t*-test.

In order to connect our observation that FAK and myosin II play a role in controlling MS-mediated integrin forces with the observation that MS creates more and/or larger internal focal adhesions with phosphorylated paxillin on Y118, we assessed focal adhesions in N27-A cells with the above inhibitors in the presence and absence of MS. N27-A cells were fixed and stained for paxillin and pY118 paxillin and are shown in **Figure 8(a)** and **Supplemental Figures 7-10**. Blocking FAK and myosin using defactinib and blebbistatin, respectively resulted in little change in focal adhesion area (**Figure 8(b)**) or paxillin (**Figure 8(c)**), but less paxillin phosphorylation (**Figure 8(d)**), when examining either total or external focal adhesions. The internal focal adhesion is so low under no MS stimulation, that neither of these treatments produced a decrease. Omecamtiv, did not show an increase in any of focal adhesion measurements when examining either the total or external focal adhesion. However, it did produce large changes in focal adhesion area, paxillin and pY118 paxillin in internal adhesions. This response matched very closely the TMS response, which also showed increases in internal adhesions. Likely, MS and omecamtiv are activating similar pathways. Indeed, MS and omecamtiv in neuronal cells both seem to activate voltage-gated Na^+^ channels, which can recruit myosin II. Under MS, both defactinib and blebbistatin diminish internal focal adhesion signatures, further supporting the hypothesis that an upstream signal like voltage-gated Na^+^ channels, activates both FAK and myosin II and is shared by both MS and omecamtiv. Omecamtiv, in addition to MS, does not enhance focal adhesion measurements over omecamtiv or MS alone, suggesting a saturating response and supporting the observation that both activate similar pathways that lead to a saturation of the ITS signal. Consequently, the response of internal focal adhesion signatures and in particular, phosphorylated paxillin seem to track well with integrin forces assessed through ITS fluorescence, placing both FAK and myosin II upstream of internal phosphorylated paxillin focal adhesions and integrin forces.

**Figure 8.**
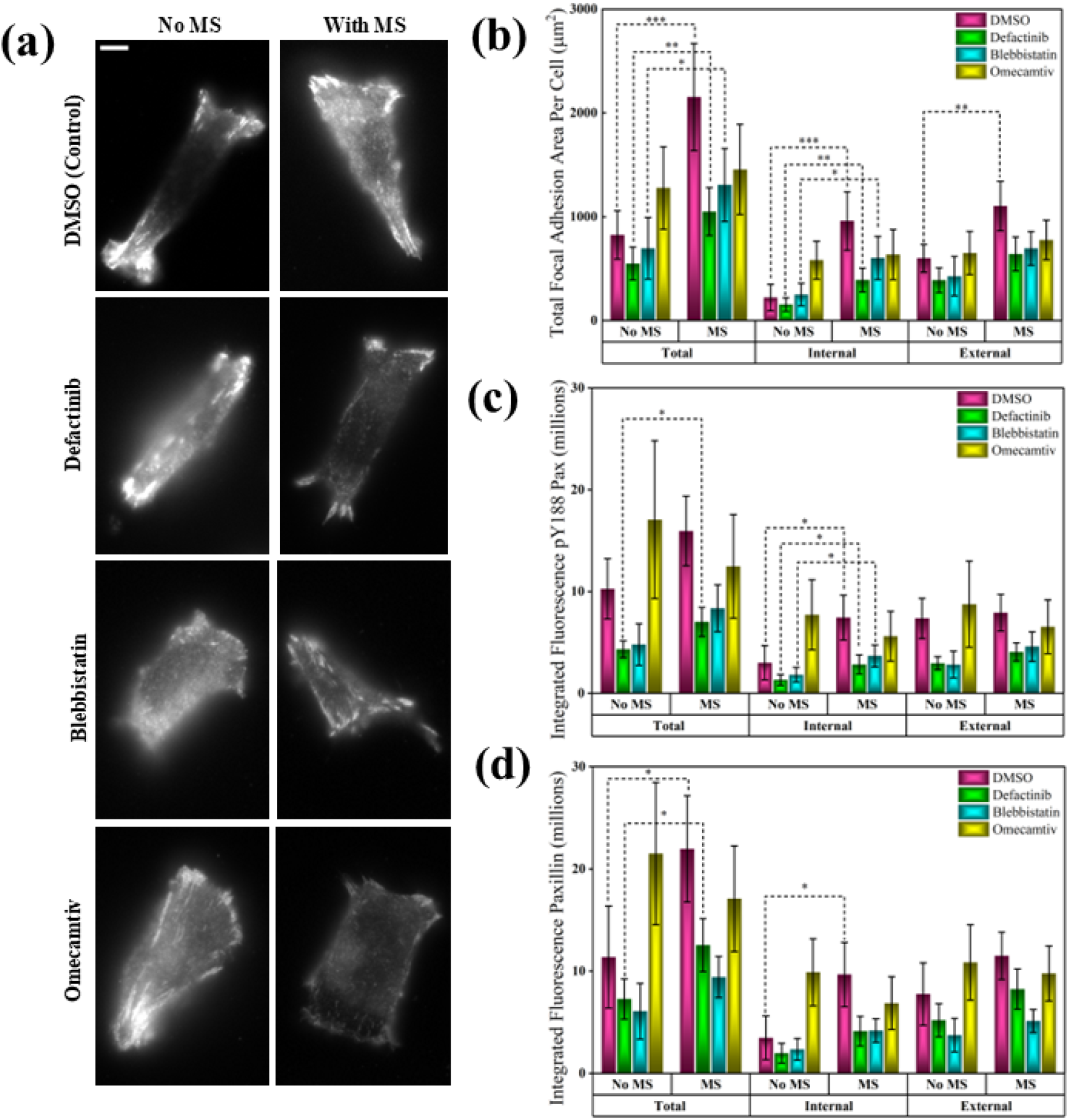
(a) Fluorescence images of N27-A cells on fibronectin coated coverslips stained for paxillin with and without MS treatment, and with either DMSO (control), 10 μM defactinib, 10 μM blebbistatin, or 10 μM omecamtiv mecarbil. Calibration bar length = 5 μm. (b) Total focal adhesion area per cell in the internal, external and total cell regions. (c) Integrated fluorescence per cell of paxillin in focal adhesions in the internal, external and total cell regions. (d) Integrated fluorescence per cell pY118 paxillin in the focal adhesions in the internal, external and total focal adhesions. (N_days_ = 2, N_total experiments_ = 2, N_cells per experiment_ = 10) Error bars are 95% confidence intervals. Asterisks * and ** and *** indicate statistically significant differences with p > 0.01, 0.001 and 0.0001 respectively using two-tailed student’s t-test. Only MS vs no MS for each inhibitor were statistically compared.

## 4. Conclusions

In this paper, the mechanosensitive response to MS has been studied in N27-A cells, a widely used *in vitro* model for PD. This behavior was examined under MS therapeutic treatment conditions that are clinically relevant for the treatment of PD. The application of MS significantly increases integrin force as measured by ITS rupture on both short (minutes) and long (hours) timescales. This enhancement was linearly dependent on magnetic field dose and non-linearly dependent on magnetic pulse repetition frequency, showing dramatic drops in response for low frequencies. Much of the enhancement by MS occurs for ITS requiring 12 pN integrin forces for rupture as compared to 54 pN integrin forces. MS treatment resulted in migration that occurred 2-4 hour after MS treatment. Increases in integrin force were attributed to both increases in integrin tension or loading rate as well as increases in integrin dynamics as ITS fluorescence signal increased after MS treatment during both the non-migratory and migratory stages. Blocking FAK or myosin II resulted in diminished integrin forces comparing cells treated with MS to those that were not, suggesting that both FAK and myosin II are involved in MS-mediated integrin forces. Commensurate with increased integrin force, MS produced more and larger internal focal adhesions that were enriched with phosphorylated paxillin on Y118, a site phosphorylated by FAK and dependent on myosin II activity. Internal focal adhesion signatures that were elicited by MS were subsequently blocked by inhibiting FAK and myosin II. Interestingly, omecamtiv, which is a cardiac-specific myosin II inhibitor seemed to act similarly to MS, increasing integrin force and internal focal adhesions. However, this drug does activate voltage-gated Na^+^ channels, suggesting that voltage-gated Na^+^ channels may be upstream of FAK and myosin II in response to MS. FAK-paxillin signaling is known to be important in dendritic spine assembly and maturation, suggesting that this MS may upregulate neuroplasticity through this mechanobiological pathway. Mechanistically detailing the response of neuronal cells to MS is lacking, particularly with respect to mechanosensitive pathways. This work suggests significant mechanosensitive pathways are activated by MS alongside the traditional secretory pathways, expanding our understanding of how MS might be used to treat different neuronal diseases like PD.

## Acknowledgements

This research was funded in part by NSF Award ECCS# 2204447.

