## Supplemental Figures for "Myosin II-FAK-paxillin Signaling Governs Mechanosensitive Responses of Neuronal Cells to Magnetic Stimulation"

and Long Que[1, *]

[1]Department of Electrical and Computer Engineering,

[2]Department of Chemical and Biological Engineering,

[3]Molecular, Cellular and Developmental Biology Program

[4]Department of Genetics, Development and Cell Biology

Iowa State University, Ames, IA, 50011, USA

[+]Equal contribution

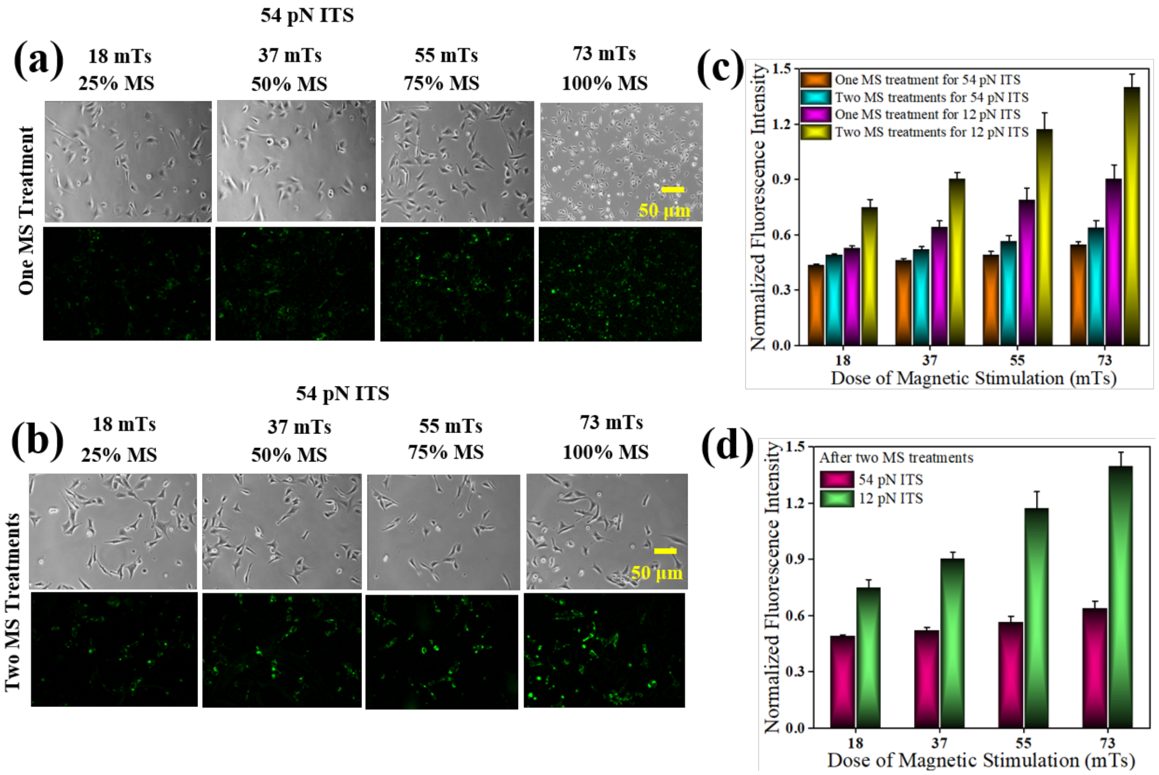


**Fig. S1**. (a) Representative phase contrast images of N27-A cells with 54 pN ITS (top row) and the corresponding fluorescence images (bottom row) subjected to one MS treatment of varied strength; (b) Representative phase contrast images of N27-A cells with 54 pN ITS (top row) and the corresponding fluorescence image (bottom row) subjected to two MS treatments of varied strength; (c) Normalized fluorescence intensity of the control sample and the samples subjected to one MS treatment and two MS treatments on either 12 pN ITS or 54 pN ITS; (d) Net changes in fluorescence intensity after subtraction of the control sample values. $N_{days} = 3$, $N_{total\ experiments} = 3$, $N_{\text{images per experiment}} = 3$.

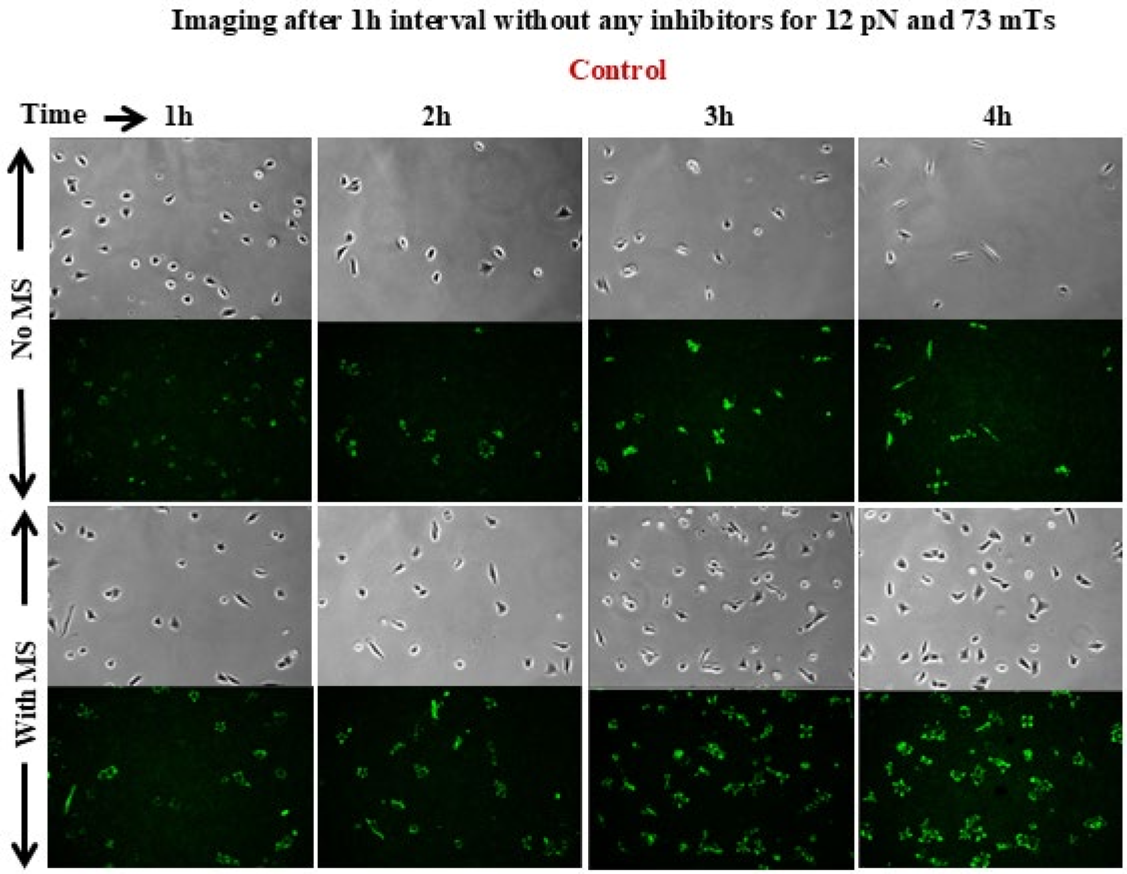


**Fig. S2**. Phase contrast and fluorescence images of N27-A cells with 12 pN ITSs without inhibitor treatment, with or without MS treatment of 73 mTs.

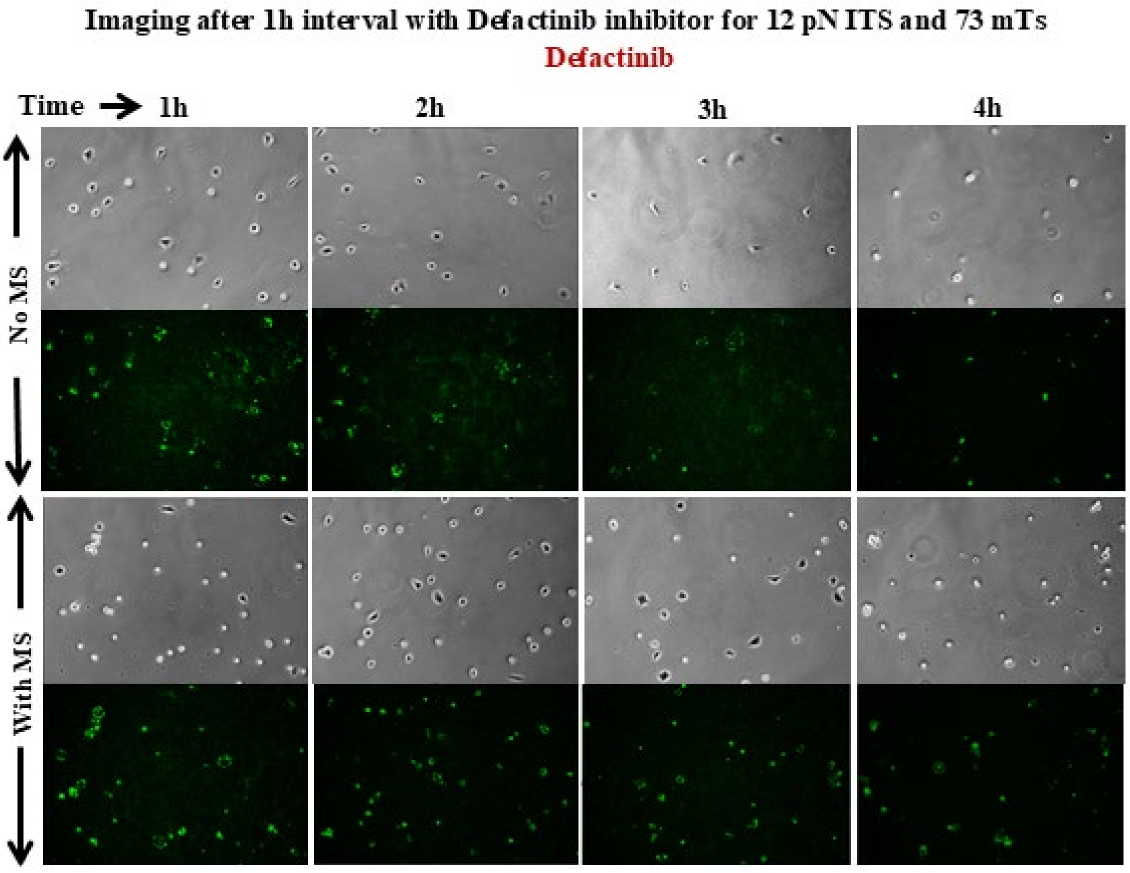


**Fig. S3**. Phase contrast and fluorescence images of N27-A cells with 12 pN ITSs with 10 μM defactinib treatment, with or without MS treatment of 73 mTs.

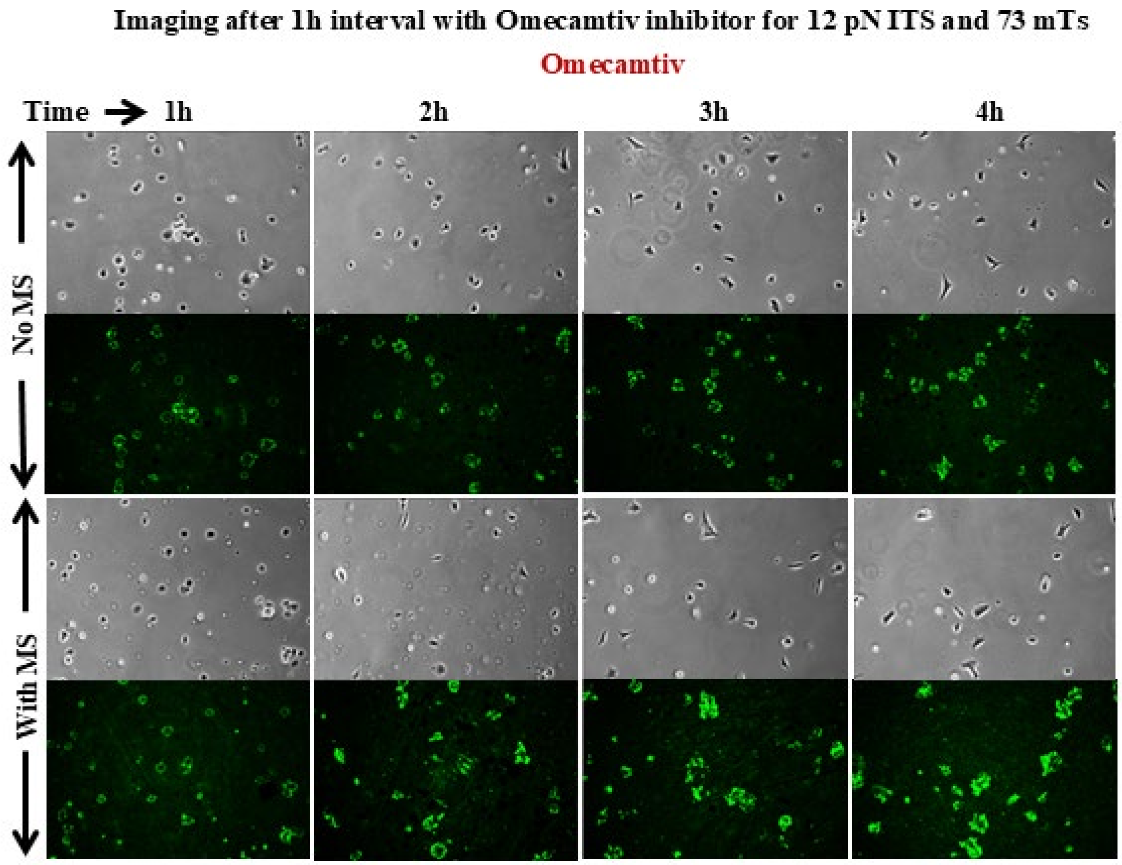


**Fig. S4**. Phase contrast and fluorescence images of N27-A cells with 12 pN ITSs with 10 μM omecamtiv treatment, with or without MS treatment of 73 mTs.

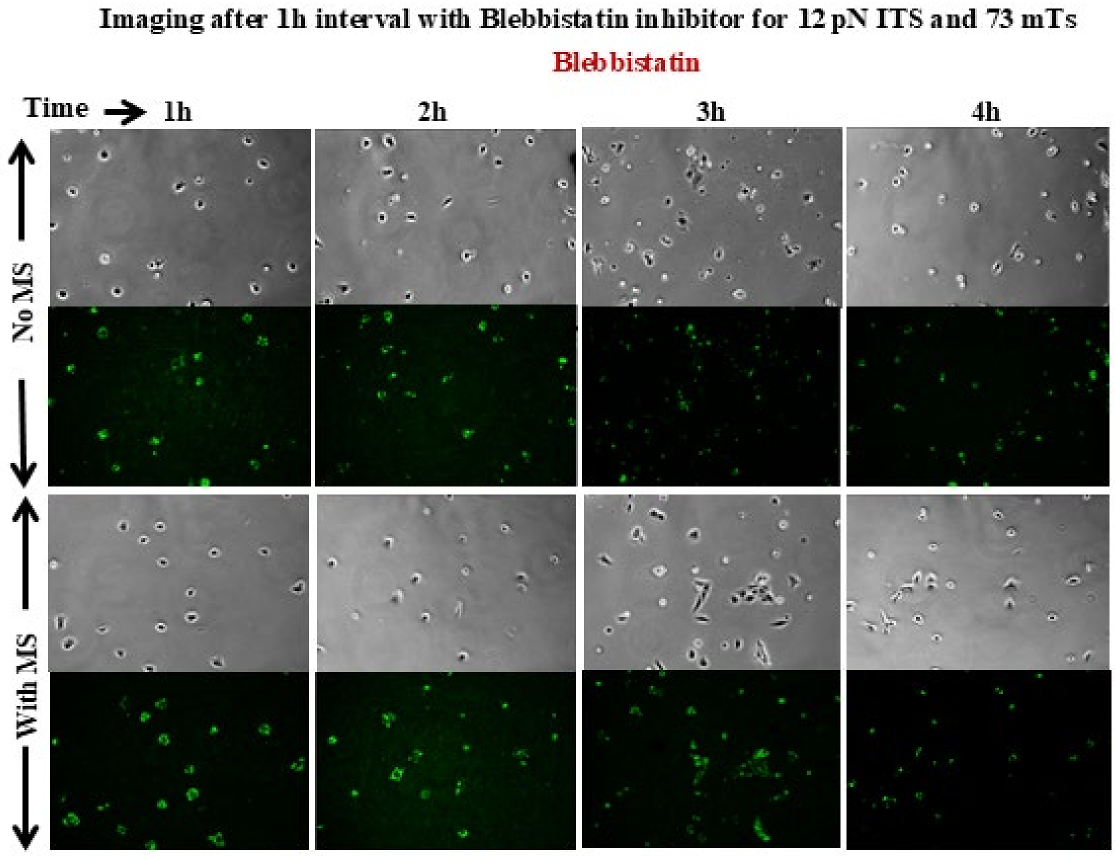


**Fig. S5**. Phase contrast and fluorescence images of N27-A cells with 12 pN ITSs with 10 μM blebbistatin treatment, with or without MS treatment of 73 mTs.

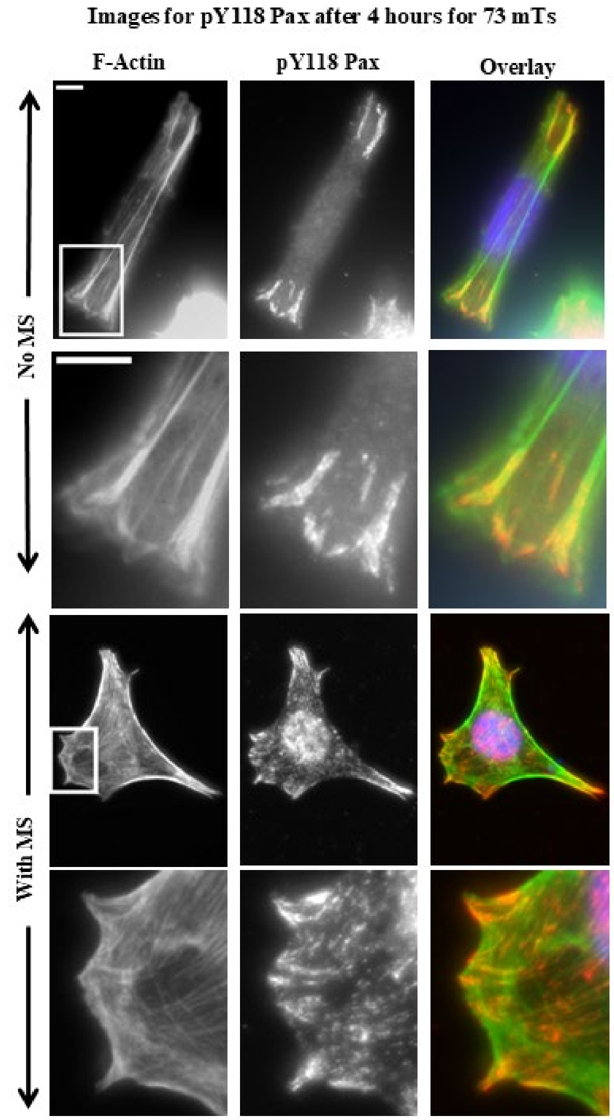


**Fig. S6.** Fluorescence images of N27-A cells on fibronectin coated coverslips stained for F-actin and pY118 paxillin with and without MS treatment. Overlays of F-actin (green), pY118 paxillin (red) and DAPI (blue) are included. Calibration bar length = 5 μm.

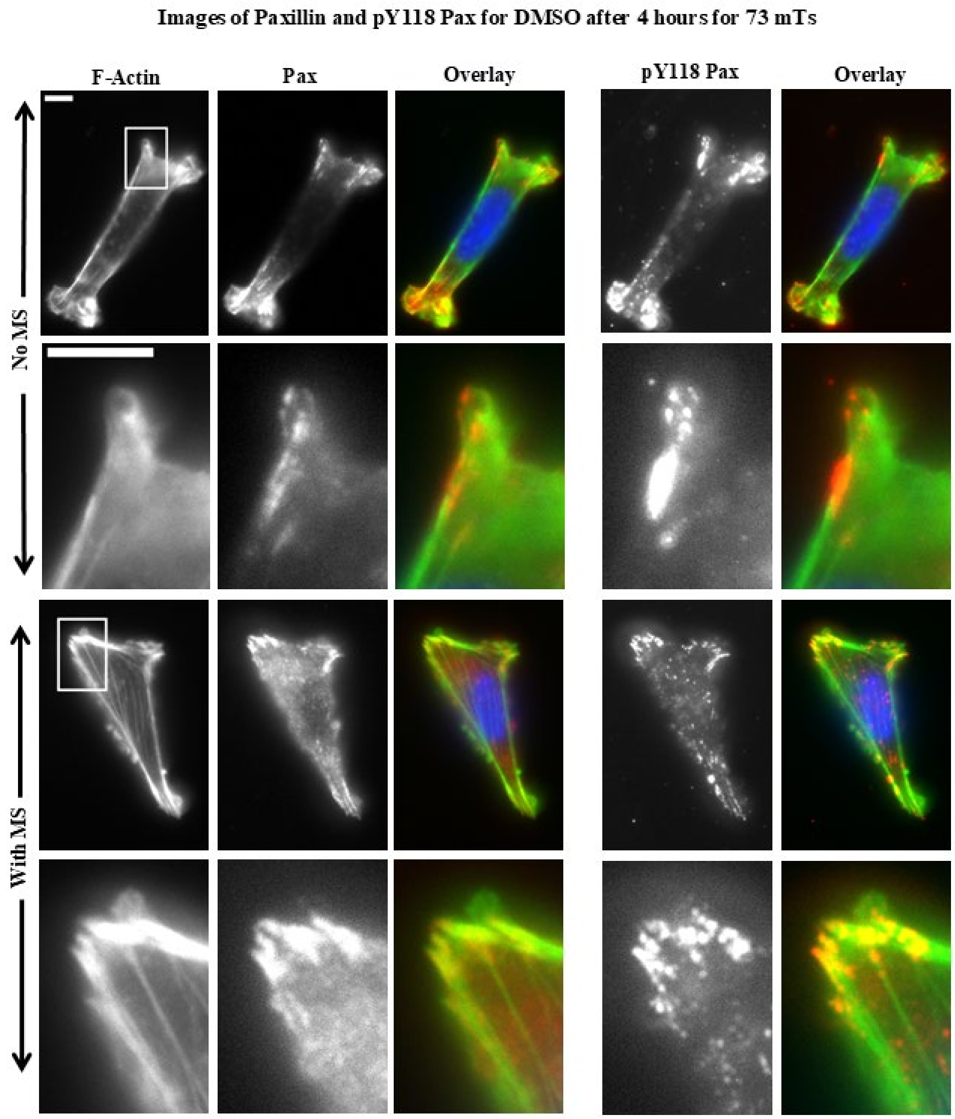


**Fig. S7**. Fluorescence images of DMSO (control) treated N27-A cells on fibronectin coated coverslips stained for F-actin, paxillin and pY118 paxillin with and without MS treatment. Overlays of F-actin (green), paxillin or pY118 paxillin (red) and DAPI (blue) are included. Calibration bar length = 5 μm.

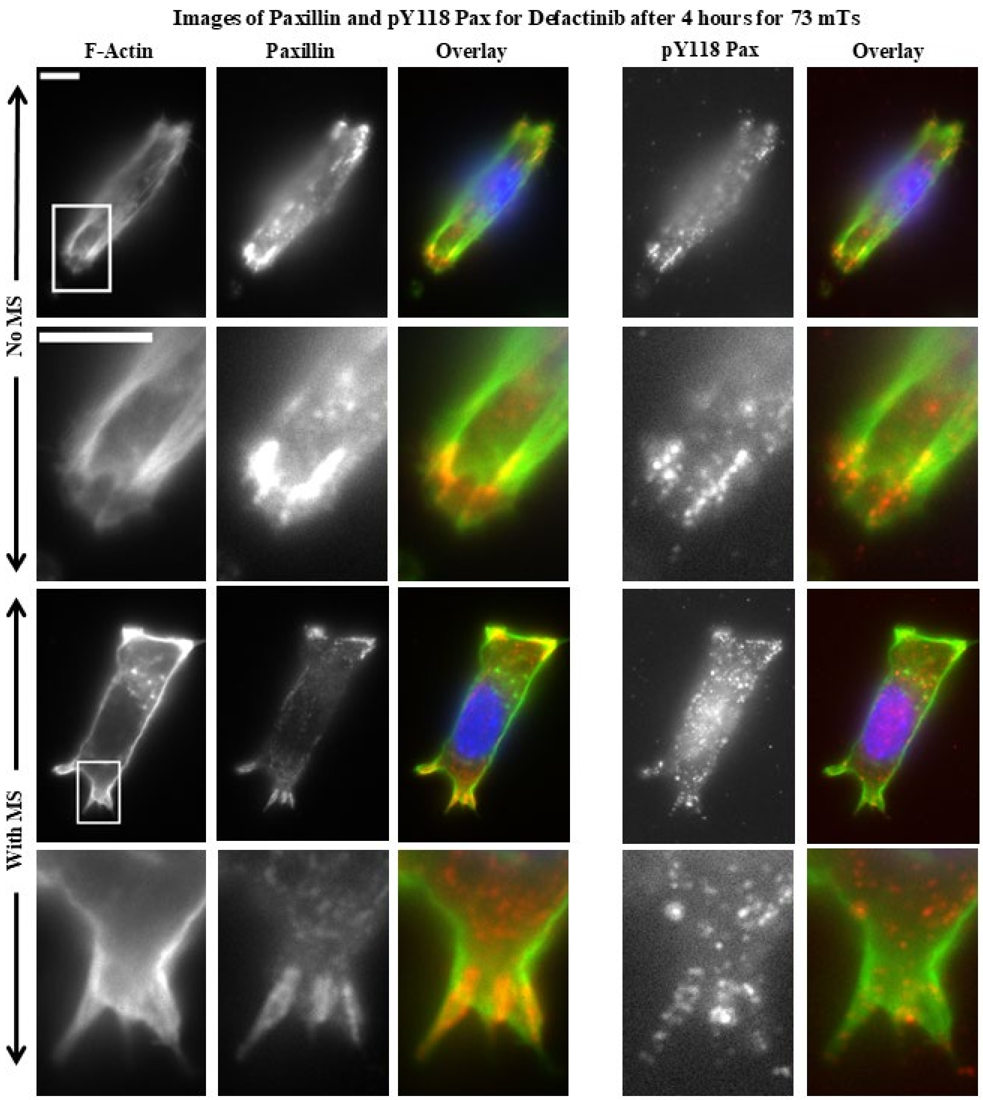


**Fig. S8**. Fluorescence images of 10 μM defactinib treated N27-A cells on fibronectin coated coverslips stained for F-actin, paxillin and pY118 paxillin with and without MS treatment. Overlays of F-actin (green), paxillin (red) or pY118 paxillin (blue) and DAPI are included. Calibration bar length = 5 μm.

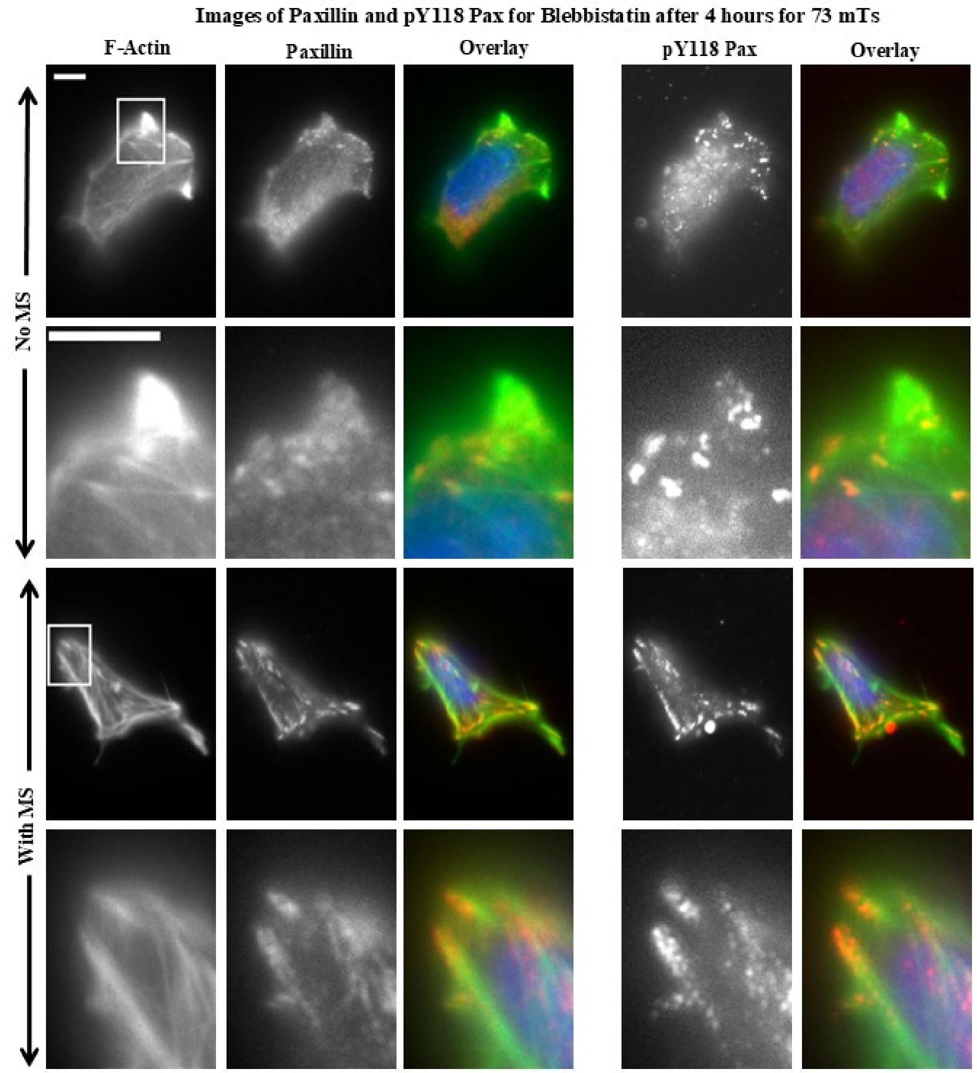


**Fig. S9**. Fluorescence images of 10 μM blebbistatin treated N27-A cells on fibronectin coated coverslips stained for F-actin, paxillin and pY118 paxillin with and without MS treatment. Overlays of F-actin (green), paxillin or pY118 paxillin (red) and DAPI (blue) are included. Calibration bar length = 5 μm.

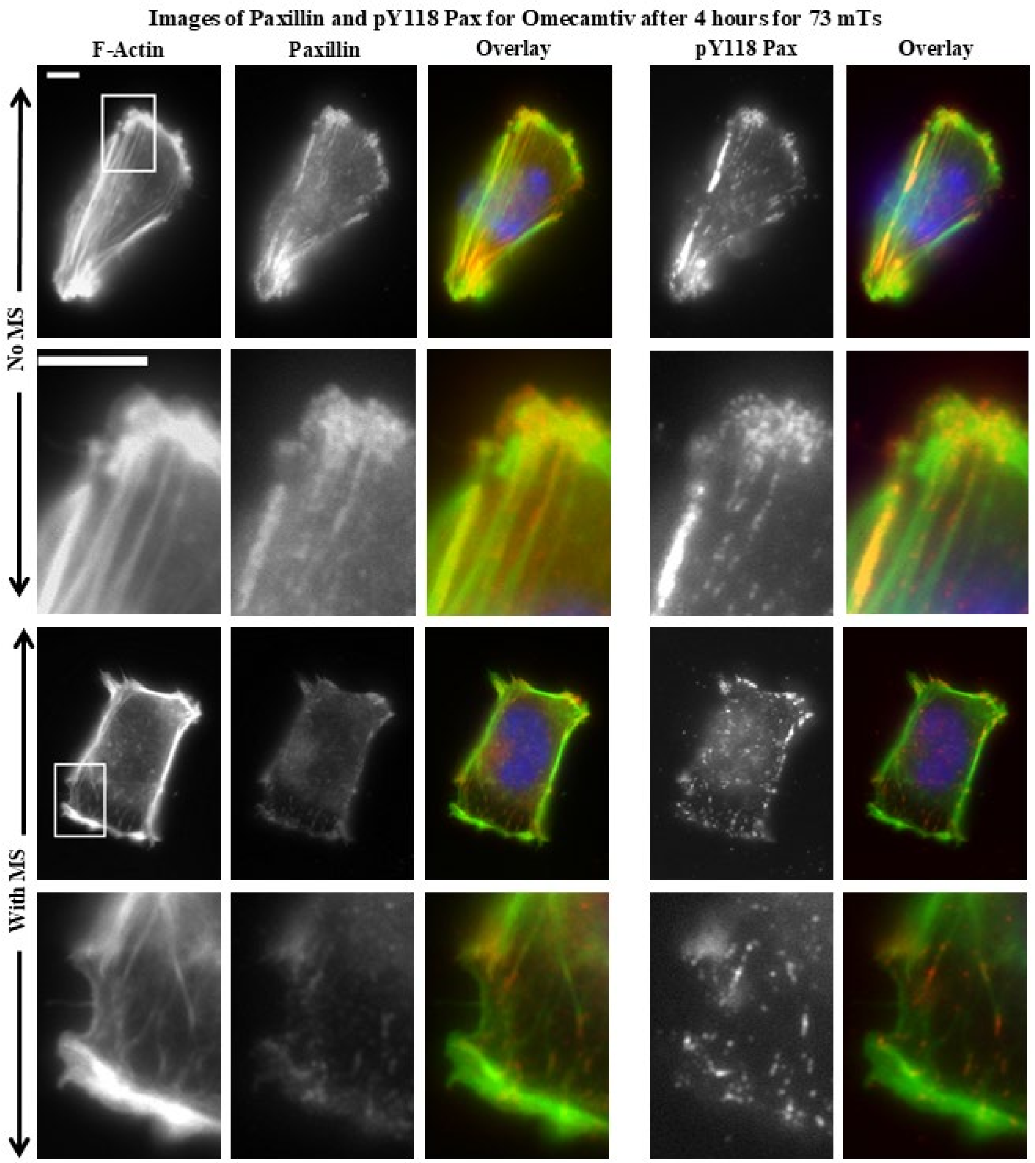


**Fig. S10**. Fluorescence images of 10 μM omecamtiv mecarbil treated N27-A cells on fibronectin coated coverslips stained for F-actin, paxillin and pY118 paxillin with and without MS treatment. Overlays of F-actin (green), paxillin or pY118 paxillin (red) and DAPI (blue) are included. Calibration bar length = 5 μm.
